# Improved specificity and efficiency of *in vivo* adenine base editing therapies with hybrid guide RNAs

**DOI:** 10.1101/2024.04.22.590531

**Authors:** Madelynn N. Whittaker, Lauren C. Testa, Dominique L. Brooks, Aidan Quigley, Hooda Said, Garima Dwivedi, Ishaan Jindal, Daphne Volpp, Ping Qu, Josh Zhiyong Wang, Michael A. Levine, Rebecca C. Ahrens-Nicklas, Qiaoli Li, Kiran Musunuru, Mohamad-Gabriel Alameh, William H. Peranteau, Xiao Wang

## Abstract

Phenylketonuria (PKU), pseudoxanthoma elasticum (PXE), and hereditary tyrosinemia type 1 (HT1) are autosomal recessive disorders linked to the *PAH*, *ABCC6*, and *FAH* and *HPD* genes, respectively. Here we evaluated the off-target editing profiles of clinical lead guide RNAs (gRNAs) that, when combined with adenine base editors (ABEs), correct the recurrent *PAH* P281L variant, *PAH* R408W variant, or *ABCC6* R1164X variant or disrupt either of two sites in the *HPD* gene (a modifier gene of HT1) in human hepatocytes. To mitigate off-target mutagenesis, we systematically screened hybrid gRNAs with DNA nucleotide substitutions. Comprehensive and variant-aware specificity profiling of these hybrid gRNAs revealed dramatically reduced off-target editing and reduced bystander editing in cells. In humanized *PAH* P281L and *ABCC6* R1164X mouse models of PKU and PXE, we showed that when formulated in lipid nanoparticles (LNPs) with ABE mRNA, selected hybrid gRNAs reverted disease phenotypes, reduced off-target editing, increased on-target editing, and reduced bystander editing *in vivo*. These studies highlight the utility of hybrid gRNAs to improve the safety and efficiency of adenine base editing therapies.

## Introduction

*In vivo* CRISPR base editing is an emerging therapeutic approach to efficiently and precisely make genomic alterations, including the direct correction of pathogenic variants, in a patient’s body^1,2^. Early results from a clinical trial of base editing to inactivate the *PCSK9* gene in patients with familial hypercholesterolemia have provided a strong proof of concept of the efficacy of adenine base editing in the human liver *in vivo*^3^. These results motivate the development of liver-centered base-editing therapies to address a wide variety of genetic disorders. In previous studies, we have demonstrated the ability of *in vivo* base editing to definitively address two such disorders in postnatal or prenatal mouse models: correction of the recurrent *PAH* P281L variant or *PAH* R408W variant in humanized mouse models of PKU, a metabolic disorder caused by increased blood phenylalanine (Phe) levels resulting from a defective phenylalanine hydroxylase (PAH) enzyme^4,5^; and inactivation of the mouse ortholog of the *HPD* gene in a mouse of model of HT1, a metabolic disorder caused by toxic metabolites resulting from a defective fumarylacetoacetate hydroxylase (FAH) enzyme, addressable by blocking the upstream 4-hydroxyphenylpyruvate dioxygenase (HPD) enzyme^6^. PXE is a connective tissue disorder marked by mineralization in various body tissues, associated with decreased blood pyrophosphate levels resulting from a defective ATP binding cassette subfamily C member 6 (ABCC6) transporter most highly expressed in the liver; the *ABCC6* R1164X variant is one of the most recurrent variants identified in PXE patients^7^.

The eighth-generation ABE, ABE8.8-m (ABE8.8), has now been validated in preclinical non-human primate studies and in the aforementioned clinical trial.^3,8,9^ As with other ABEs, it can use its adenosine deaminase domain to efficiently catalyze A⟶G edits on the sense or antisense strands of genes and correct a substantial proportion of human pathogenic variants. ABE8.8 also has among the narrowest editing windows of the eighth-generation ABEs^10^. If the position of the target adenine base lies within the window of ABE8.8, a narrow window limits the potential for two undesirable consequences of base editing: bystander editing and off-target editing. Bystander editing of one or more adenine bases near the target adenine base can be counterproductive if it introduces new pathogenic variants alongside the desired corrective editing of the original pathogenic variant. Off-target editing, a general concern with any type of CRISPR editing, entails mutagenesis at genomic sites other than the desired on-target site. Besides the use of a narrow-window ABE, strategies that can simultaneously reduce off-target editing and bystander editing would be of substantial value in improving the safety and efficiency of ABE therapies.

In this work, we evaluated clinical lead gRNAs that, in combination with an ABE, achieve therapeutically desirable edits in *PAH*, *ABCC6*, or *HPD* in hepatocytes. To reduce the off-target editing by these gRNAs, we explored a previously reported strategy, the *in vitro* or *in cellulo* use of hybrid gRNAs in which certain positions of the spacer sequence are substituted with DNA nucleotides^11–17—a^ strategy compatible with the use of mRNA-LNPs for *in vivo* delivery. In systematically assessing the effects of a series of hybrid gRNAs on the off-target profiles of clinical lead gRNAs, we unexpectedly observed that hybrid gRNAs have the potential to reduce bystander editing as well. Importantly, we found that hybrid gRNAs that reduced off-target editing by the lead *PAH* P281L-correcting and *ABCC6* R1164X-correcting ABE/gRNA combinations *in vivo*, in PKU and PXE humanized mouse models, did so without compromising *in vivo* editing efficiency. On the contrary, the hybrid gRNAs significantly increased the desired corrective editing in the liver while simultaneously reducing unwanted bystander editing.

## Results

### Optimizing correction of the *PAH* P281L variant *in cellulo*

In a previous study, we characterized a clinical lead gRNA (herein termed “PAH1”) targeting the human-specific protospacer sequence TCAC**<u>A</u>**GTTCGGGGGTATACA, with the corresponding protospacer-adjacent motif (PAM) TGG, to correct the *PAH* P281L variant^4^ (**Fig. 1a**). The variant adenine base lies in position 5 of the protospacer sequence. A potential bystander adenine base lies nearby in position 3; bystander A⟶G editing would result in a splice site disruption that has been reported to be pathogenic for PKU^18^ and thus would be undesirable. In combination with ABE8.8, the PAH1 gRNA results in efficient corrective editing of the P281L variant *in cellulo* (in HuH-7 hepatocytes homozygous for the P281L variant, derived from the HuH-7 human hepatoma cell line) and *in vivo* (in mouse liver), albeit with sizeable levels of bystander editing^4^.

**Fig. 1.**
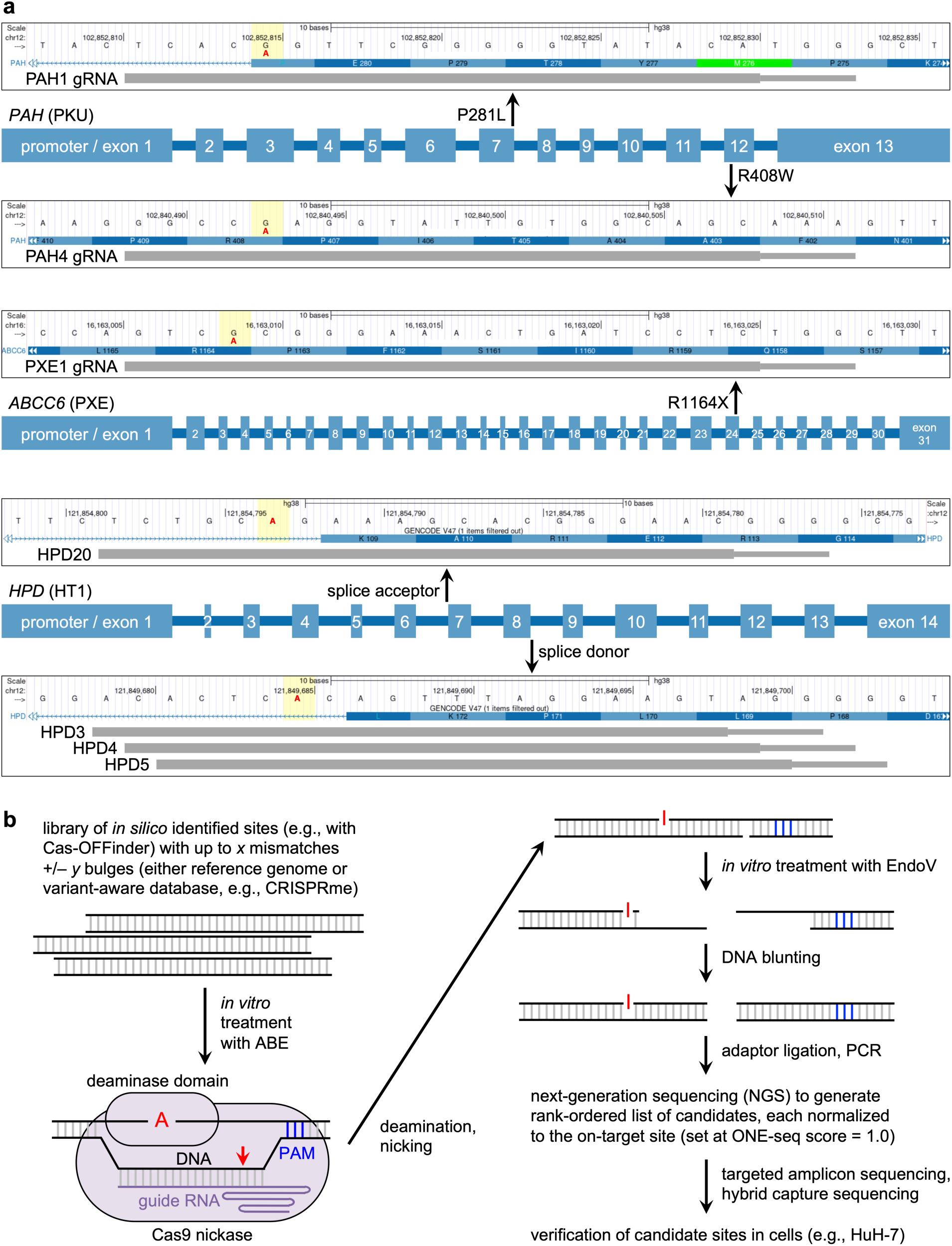
Assessment of on-target and off-target editing by candidate clinical lead guide RNAs. **a**, Schematics of the genomic sites of the five loci that are evaluated in this study, adapted from the UCSC Genome Browser (GRCh38/hg38). Each vertical yellow bar indicates either the G altered to A (in red) by a pathogenic variant, or the target A (in red) at a splice site intended for disruption. Each horizontal grey bar indicates the protospacer (thick) and protospacer-adjacent motif (PAM) (thin) sequences targeted by the respective guide RNA (gRNA). **b**, Schematic of the ONE-seq methodology.

Most CRISPR-based off-target assay techniques are designed to detect double-strand breaks in DNA, whether *in vitro* or *in cellulo*. However, base editors like ABE8.8 do not directly make double-strand breaks, and so conventional off-target assays such as GUIDE-seq do not accurately reflect off-target editing by base editors. We therefore utilized an ABE-tailored version of OligoNucleotide Enrichment and sequencing (ONE-seq) to nominate off-target sites^8,19^, followed by genomic sequencing to verify these candidate sites (**Fig. 1b**). ONE-seq is a biochemical assay that uses a synthetic genomic library, selected by similarity between the protospacer DNA sequence targeted by the gRNA and sequences anywhere in a reference human or mouse genome, using tools like Cas-OFFinder^20^. For any given gRNA, the library can comprise tens of thousands of oligonucleotides. The library is mixed *in vitro* with a ribonucleoprotein (RNP) comprising recombinant ABE protein complexed with the synthetic gRNA. The RNP nicks certain oligonucleotide sequences on one strand and deaminates an adenine base on the other strand. EndoV is used to cleave the other strand adjacent to the deaminated base, resulting in the equivalent of a double-strand break. Next-generation sequencing (NGS) then quantifies the frequency with which each unique oligonucleotide sequence was edited *in vitro*, generating a rank-ordered list of candidate off-target sites, normalized to the on-target site, which receives a ONE-seq score of 1.0. Typically, the on-target site is at or near the top of the list. A site with a ONE-seq score of 0.01 had one-hundredth the *in vitro* editing activity of the on-target site.

For the ABE8.8/PAH1 combination, we had previously performed a ONE-seq analysis followed by targeted amplicon sequencing of ≈50 top-ranked sites that verified one genomic site of low-level off-target mutagenesis in HuH-7 hepatocytes.^4^ We now undertook a deeper analysis of the ONE-seq rank-ordered list, using hybrid capture sequencing to assess the entire set of 280 genomic sites with ONE-seq scores greater than 0.01 (**Supplementary Table 1**). We verified an additional six sites of off-target mutagenesis in ABE8.8/PAH1 mRNA/gRNA-transfected P281L HuH-7 hepatocytes (**Fig. 2a**). None of the seven sites raises a concern of oncogenic risk (see **Supplementary Note 1**). The third verified off-target site (PAH1_OT3) displayed the most off-target editing, with 1.3% editing.

**Fig. 2.**
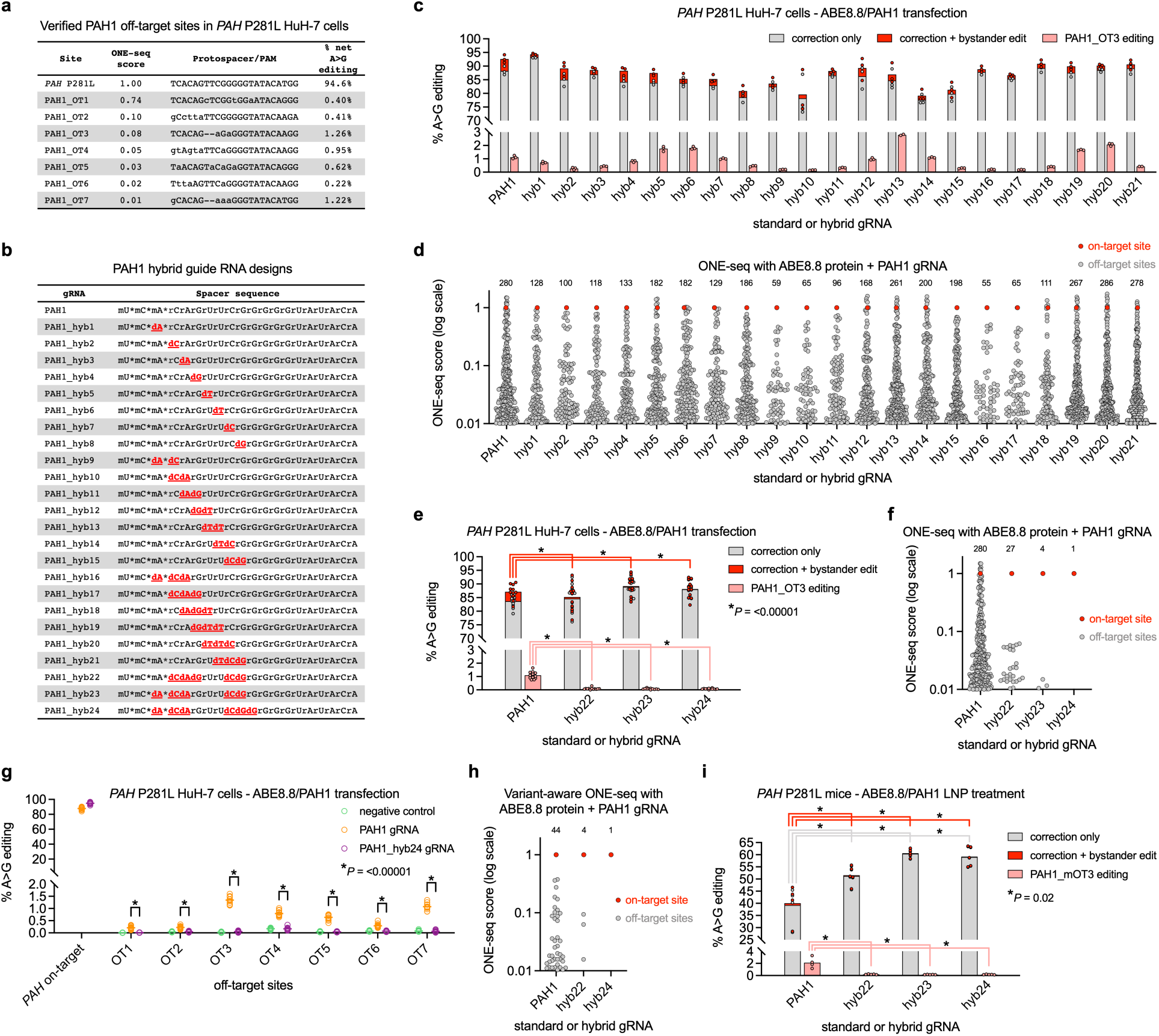
Optimizing correction of the *PAH* P281L variant *in cellulo* and *in vivo*. **a**, Hybrid capture sequencing of ONE-seq-nominated sites in treated vs. control *PAH* P281L HuH-7 cells with PAH1 gRNA in combination with ABE8.8 mRNA (*n* = 3 treated and 3 untreated biological replicates), with seven human sites verified to have off-target editing. **b**, Spacer sequences of PAH1 standard and hybrid gRNAs. “rN” = ribonucleotide; “dN” = deoxyribonucleotide; “m” = 2’-O-methylation; * = phosphorothioate linkage. DNA substitutions are in red bold and underlined. **c**, A-to-G editing of *PAH* on-target site (correction of P281L variant only, or correction plus unwanted nonsynonymous bystander editing) or verified PAH1 human off-target site (PAH1_OT3) in *PAH* P281L HuH-7 cells treated with PAH1 standard or hybrid gRNA in combination with ABE8.8 mRNA (*n* = 2–3 biological replicates per condition). **d**, Number of sites with ONE-seq score > 0.01 for each indicated PAH1 standard or hybrid gRNA with ABE8.8 protein. **e**, A-to-G editing of *PAH* on-target site or verified PAH1 human off-target site in *PAH* P281L HuH-7 cells treated with PAH1 standard or hybrid gRNA in combination with ABE8.8 mRNA (*n* = 11–12 biological replicates per condition). **f**, Number of sites with ONE-seq score > 0.01 for each indicated PAH1 standard or hybrid gRNA with ABE8.8 protein. **g**, Total A-to-G editing of *PAH* on-target site or any of seven verified PAH1 human off-target sites in untreated vs. treated *PAH* P281L HuH-7 cells with PAH1 standard or PAH1_hyb24 gRNA in combination with ABE8.8 mRNA (*n* = 11–12 biological replicates per condition). **h**, Number of variant (non-reference-genome) sites with ONE-seq score > 0.01 for each indicated PAH1 standard or hybrid gRNA with ABE8.8 protein. **i**, A-to-G editing of *PAH* on-target site or verified PAH1 mouse off-target site (PAH1_mOT3) in homozygous *PAH* P281L mice treated with 2.5 mg/kg dose of LNPs with PAH1 standard or hybrid gRNA in combination with ABE8.8 mRNA (*n* = 4–5 animals per group). Means are shown; *P* values were calculated with the Mann– Whitney *U* test.

To assess whether hybrid gRNAs could reduce the off-target editing at the verified sites, we designed a series of 21 synthetic hybrid gRNAs (“PAH1_hyb1” through “PAH1_hyb21”) with single, double, or triple DNA nucleotide substitutions ranging from positions 3 through 10 in the spacer sequence (**Fig. 2b**). We transfected ABE8.8 mRNA in combination with the PAH1 standard gRNA and each of the hybrid gRNAs into P281L HuH-7 hepatocytes and assessed for (1) on-target P281L corrective editing, (2) bystander editing at the on-target site, and (3) PAH1_OT3 off-target editing in each of the samples (**Fig. 2c**). While most of the PAH1 hybrid gRNAs had comparable on-target editing to the PAH1 standard gRNA (≈90%), several had modestly decreased editing (as low as ≈80%). Notably and unexpectedly, most of the hybrid gRNAs reduced bystander editing (from 4.4% with the standard gRNA to as low as ≈1%). The PAH1 hybrid gRNAs had highly varied effects on PAH1_OT3 off-target editing. Although triple DNA nucleotide substitutions generally reduced PAH1_OT3 off-target editing more than single or double substitutions, a couple of gRNAs with triple substitutions had greater PAH1_OT3 off-target editing than the standard gRNA. None of the 21 PAH1 hybrid gRNA fully eliminated

PAH1_OT3 off-target editing. As an orthogonal method of off-target profiling, we performed ONE-seq with ABE8.8 and each of the 21 PAH1 hybrid gRNAs (**Fig. 2d**). We used the number of genomic sites with ONE-seq scores greater than 0.01 as a metric of gRNA specificity—the fewer the sites, the less potential for off-target editing. There was good agreement between the results of the site-specific PAH1_OT3 analysis and the more general ONE-seq off-target profiling for PAH1 hybrid gRNAs.

To further reduce off-target editing, we combined triple and double substitutions within a single hybrid gRNA. We chose the triple substitutions of PAH_hyb17 (positions 4, 5, and 6) or PAH_hyb16 (positions 3, 4, and 5), which maximally reduced PAH1_OT3 off-target editing and maximally reduced bystander editing while preserving on-target editing, and added the double substitutions of PAH_hyb15 (positions 9 and 10), which substantially reduced PAH1_OT3 off-target editing and bystander editing, yielding PAH1_hyb22 and PAH1_hyb23. We added an additional substitution in position 11 to PAH1_hyb23, yielding PAH1_hyb24. We compared these three hybrid gRNAs directly against the PAH1 standard gRNA and negative controls via mRNA/gRNA transfection in P281L HuH-7 hepatocytes; all three significantly reduced PAH1_OT3 off-target editing and bystander editing (**Fig. 2e**). ONE-seq confirmed the improved off-target profiles of the three new hybrid gRNAs relative to the 21 original hybrid gRNAs; indeed, PAH1_hyb24 had zero off-target sites with ONE-seq scores greater than 0.01 (**Fig. 2f**). At all seven verified PAH1 off-target sites, OT1 through OT7, PAH1_hyb24 reduced the detectable off-target editing to the background levels of the negative controls (**Fig. 2g**).

All standard off-target assessment techniques share a critical limitation: each is tied to the specific individual genome represented by the cells or by the *in vitro* genomic DNA sample used for analysis. For this reason, most off-target analyses have simply ignored the potential for naturally occurring human genetic variation to create novel off-target editing sites in some patients. A cautionary tale is provided by the finding that the gRNA used in the recently approved CRISPR-based therapy for sickle cell disease, exa-cel, has substantial off-target editing in hematopoietic stem cells at a genomic site created by a genetic variant present in ≈10% of individuals of African ancestry, resulting in both indel mutations and chromosomal rearrangements^21^. The ONE-seq technique is uniquely capable of performing a variant-aware off-target analysis, accommodating naturally occurring human genetic variation by using oligonucleotide libraries designed not just using the reference human genome but also incorporating data from the 1000 Genomes Project, the Human Genome Diversity Project, etc.

We used the bioinformatic tool CRISPRme^21^ to design a variant-aware ONE-seq library for the PAH1 protospacer with ≈7,000 non-reference-genome oligonucleotides. In performing a variant-aware ONE-seq experiment with the ABE8.8/PAH1 combination, we identified 40 variant sites with ONE-seq scores greater than 0.01 (**Fig. 2h**). One of the top variant sites, with a ONE-seq score of 0.19, is a rare singleton 1-bp deletion that alters a reference genomic site that has 7 mismatches with the on-target protospacer, with an NTG PAM, into a site that has 3 mismatches, with an NGG PAM (**Extended Fig. 1**). Another top variant site, with a ONE-seq score of 0.11, is common enough to have been previously cataloged as rs76813758, a single-nucleotide variant that alters a reference genomic site that has 4 mismatches with the on-target protospacer, with an NGG PAM, into a site that has 3 mismatches, with an NGG PAM; the variant is present in ≈8% of individuals of African ancestry and ≈3% of individuals of European ancestry (**Extended Fig. 1**).

One limitation of variant-aware off-target analysis is that upon identifying candidate variant sites, it can be challenging to verify whether editing actually occurs at any of those sites in the therapeutically relevant cells (e.g., hepatocytes) if there is no way to obtain such cells from individuals with those variants. We reasoned that the use of hybrid gRNAs should reduce the potential for off-target editing not only at reference genomic sites but also at variant sites, mitigating the need to evaluate variant sites *in cellulo*. Accordingly, we performed variant-aware ONE-seq experiments for the ABE8.8/PAH1_hyb22 and ABE8.8/PAH1_hyb24 combinations (**Fig. 2h**, **Extended Fig. 1**). With PAH1_hyb22, there were just 3 variant sites with ONE-seq scores greater than 0.01, and with PAH1_hyb24, there were no variant sites with ONE-seq scores greater than 0.01. For rs76813758, the ONE-seq score dropped from 0.11 to 0.004 with PAH1_hyb22 and to 0.0001 with PAH1_hyb24 (one-ten thousandth of the *in vitro* editing activity of the on-target site, within the background of the assay).

### Optimizing correction of the *PAH* P281L variant *in vivo*

Although our rational exploration of hybrid gRNAs, culminating in PAH1_hyb24, was successful in rendering editing at the seven verified PAH1 off-target sites undetectable *in cellulo* while preserving on-target editing efficiency *in cellulo*, it remained to be answered whether hybrid gRNAs could function as effectively *in vivo*. We formulated LNPs with ABE8.8 mRNA and PAH1 standard gRNA, PAH1_hyb22 gRNA, PAH1_hyb23, or PAH1_hyb24, exactly paralleling LNPs we used in a recently published study^4^. We administered the LNPs to homozygous humanized P281L PKU mice at a dose of 2.5 mg/kg. We observed that only the PAH1_hyb23 and PAH1_hyb24 gRNAs resulted in complete normalization of blood Phe levels (mean <125 µmol/L) by 48 hours after treatment, significantly outperforming the PAH1 standard gRNA (**Extended Fig. 2**). With ABE8.8/PAH1 LNPs, there was mean 40% whole-liver P281L corrective editing; with each of the hybrid gRNAs, there was mean 50%–60% editing, establishing that the use of the hybrid gRNAs did not compromise and in fact significantly improved on-target editing (**Fig. 2i**). The increased on-target editing was accompanied by significantly reduced bystander editing (mean 0.8% editing with standard gRNA compared to mean 0.2%–0.3% editing with hybrid gRNAs).

We performed ONE-seq for the ABE8.8/PAH1 combination against the reference mouse genome (**Supplementary Table 2**). In interrogating the candidate sites with the top ONE-seq scores in liver samples from the LNP-treated mice, we verified a site (PAH1_mOT3) with mean 2.1% off-target editing with the PAH1 standard gRNA (**Fig. 2i**). PAH1_hyb22 gRNA, PAH1_hyb23, and PAH1_hyb24 all significantly reduced the off-target editing at this site (<0.2%).

### Optimizing correction of the *ABCC6* R1164X variant *in cellulo*

Out of the recurrent *ABCC6* variants causative of PXE, we focused on the R1164X variant because of features suggesting it would be amenable to adenine base editing (**Fig. 1a**, **Fig. 3a**), namely the positioning vis-à-vis a TGG PAM compatible with a gRNA (designated PXE1) with the protospacer sequence GTC**<u>A</u>**CGGGAAACTGATCCTC, the variant adenine base lying in position 4, and potential bystander adenine bases lying in positions 9, 10, and 11, outside the reported editing window of ABE8.8. We used prime editing to generate a homozygous R1164X HuH-7 cell line (**Supplementary Note 2**). Transfection of the cell line with ABE8.8 mRNA and the PXE1 gRNA achieved substantial corrective editing of the R1164X variant (32%), albeit with a moderate amount of unwanted bystander editing (6.3%) (**Fig. 3b**). For the ABE8.8/PXE1 combination, we performed a ONE-seq analysis followed by targeted amplicon sequencing of top-ranked sites that verified six genomic sites of off-target mutagenesis in the HuH-7 cells (**Supplementary Table 3**). None of the six sites raises a concern of oncogenic risk (see **Supplementary Note 1**). The first and second verified off-target sites (PXE1_OT1, PXE1_OT2) displayed the most off-target editing.

**Fig. 3.**
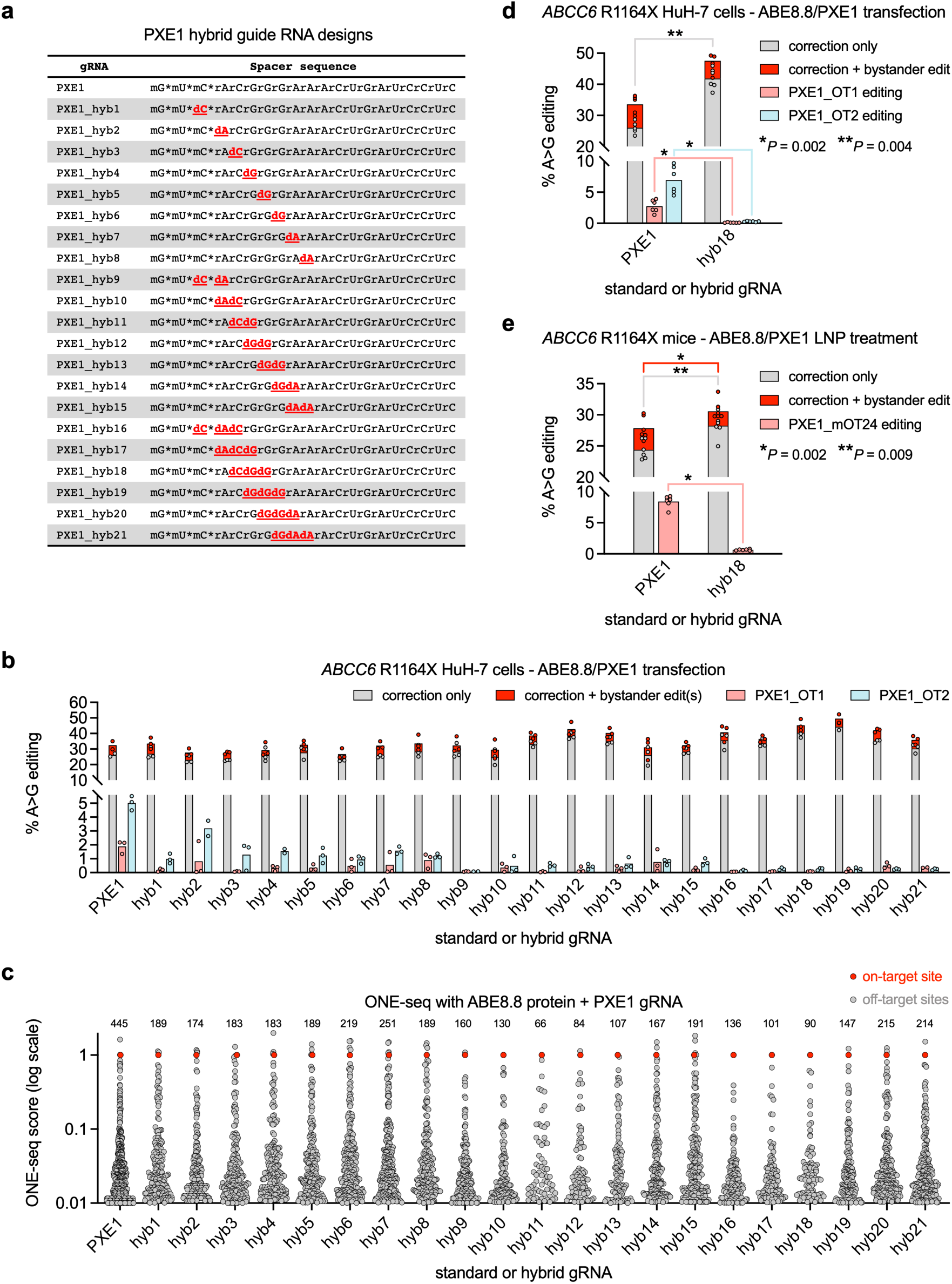
Optimizing correction of the *ABCC6* R1164X variant *in cellulo* and *in vivo*. **a**, Spacer sequences of PXE1 standard and hybrid gRNAs. “rN” = ribonucleotide; “dN” = deoxyribonucleotide; “m” = 2’-O-methylation; * = phosphorothioate linkage. DNA substitutions are in red bold and underlined. **b**, A-to-G editing of *ABCC6* on-target site (correction of R1164X variant only, or correction plus unwanted nonsynonymous bystander editing) or either of two verified PXE1 human off-target sites (PXE1_OT1, PXE_OT2) in *ABCC6* R1164X HuH-7 cells treated with PXE1 standard or hybrid gRNA in combination with ABE8.8 mRNA (*n* = 2–3 biological replicates per condition). **c**, Number of sites with ONE-seq score > 0.01 for each indicated PXE1 standard or hybrid gRNA with ABE8.8 protein. **d**, A-to-G editing of *ABCC6* on-target site or either of two verified PXE1 human off-target sites in *ABCC6* R1164X HuH-7 cells treated with PXE1 standard or PXE1_hyb18 gRNA in combination with ABE8.8 mRNA (*n* = 5– 6 biological replicates per condition). **e**, A-to-G editing of *ABCC6* on-target site or verified PXE1 mouse off-target site (PXE1_mOT24) in homozygous *ABCC6* R1164X mice treated with 2.5 mg/kg dose of LNPs with PXE1 standard or PXE1_hyb18 gRNA in combination with ABE8.8 mRNA (*n* = 6 animals per group). Means are shown; *P* values were calculated with the Mann–Whitney *U* test.

As we had done with the PAH1 gRNA, we designed a series of 21 hybrid gRNAs for PXE1 (PXE1_hyb1 through PXE1_hyb21) with single, double, or triple DNA nucleotide substitutions ranging from positions 3 through 10 in the spacer sequence (**Fig. 3a**), matching the designs of the PAH1 hybrid gRNAs. Upon transfection of ABE8.8 mRNA and each of the gRNAs into R1164X HuH-7 hepatocytes, we found that all of the hybrid gRNAs reduced off-target editing at the OT1 and OT2 sites, some of the hybrid gRNAs increased on-target editing (to as high as ≈50%), and some of the hybrid gRNAs reduced bystander editing (to as low as ≈3%) (**Fig. 3b**). We performed ONE-seq for each of the hybrid gRNAs and, judging by the number of genomic sites with ONE-seq scores greater than 0.01, improvement of off-target editing was concordant with the effects on the individual OT1 and OT2 sites (**Fig. 3c**). We further evaluated PXE1_hyb18 and found that it significantly increased on-target editing and significantly, though not entirely, reduced off-target editing at the OT1 and OT2 sites (**Fig. 3d**). It is possible that additional DNA nucleotide substitutions would yield hybrid gRNAs that further reduce off-target editing.

### Optimizing correction of the *ABCC6* R1164X variant *in vivo*

We used CRISPR-Cas9 targeting in mouse embryos to generate a humanized PXE model, in the C57BL/6J background, in which we replaced a small portion of the endogenous mouse *Abcc6* exon 24 with the orthologous human sequence spanning the PXE1 protospacer/PAM sequences and containing the R1164X variant (**Supplementary Note 3**). Consistent with prior studies of *Abcc6* knockout mice^7^, homozygous R1164X mice had reduced blood pyrophosphate levels compared to wild-type littermates (**Extended Fig. 3**).

We formulated LNPs with ABE8.8 mRNA and either PXE1 standard gRNA or PXE1_hyb18 gRNA. We administered the LNPs to homozygous R1164X mice at a dose of 2.5 mg/kg. With ABE8.8/PXE1 LNPs, there was mean 24% whole-liver P281L corrective editing in the absence of bystander editing, and mean 3.5% corrective editing with bystander editing; with the hybrid gRNA, 29% and 2.1% editing, respectively (**Fig. 3e**). The PXE1 hybrid gRNA significantly increased the on-target editing and reduced bystander editing *in vivo*, as was observed with PAH1 hybrid gRNAs *in vivo*. LNP treatment also largely normalized the blood pyrophosphate levels in the homozygous R1164X mice (**Extended Fig. 3a**).

We performed ONE-seq for the ABE8.8/PXE1 combination against the reference mouse genome (**Supplementary Table 4**). In interrogating the candidate sites with the top ONE-seq scores in liver samples from the LNP-treated mice, we verified a site (PXE1_mOT24) with mean 8.4% off-target editing with the PXE1 standard gRNA (**Fig. 3e**). The PXE1_hyb18 gRNA significantly reduced the off-target editing at this site, with mean 0.6% editing. We also assessed blood alanine aminotransferase (ALT) and cytokine/chemokine levels in LNP-treated mice, and we observed similar safety profiles with the standard and hybrid gRNAs (**Extended Fig. 3b**, **Supplementary Table 5**).

### Optimizing targeting of two distinct sites in the *HPD* gene with hybrid gRNAs

Lacking any previous data regarding adenine base editing to inactivate the *HPD* gene—our previous study assessed only cytosine base editing—we screened for precise disruption of a splice donor or acceptor site in the human *HPD* gene by adenine base editing (**Supplementary Note 4**). We observed the highest levels of editing with the gRNAs designated “HPD20”, “HPD3”, “HPD4”, and “HPD5”, targeting either the splice acceptor of exon 7 or the splice donor of exon 8 (**Fig. 1a**, **Extended Fig. 4a**). To evaluate for off-target editing, we used two ONE-seq libraries against the reference human genome: a combined library for the cluster of closely spaced HPD3, HPD4, and HPD5 protospacers (consecutive protospacers shifted by one nucleotide), and a library for the HPD20 protospacer. Targeted amplicon sequencing of top-ranked sites in ABE8.8/gRNA-treated wild-type HuH-7 hepatocytes verified one genomic site of off-target mutagenesis each for HPD3 and HPD4, and none for HPD5 and HPD20 (**Extended Fig. 4b**, **Supplementary Tables 6–9**).

**Fig. 4.**
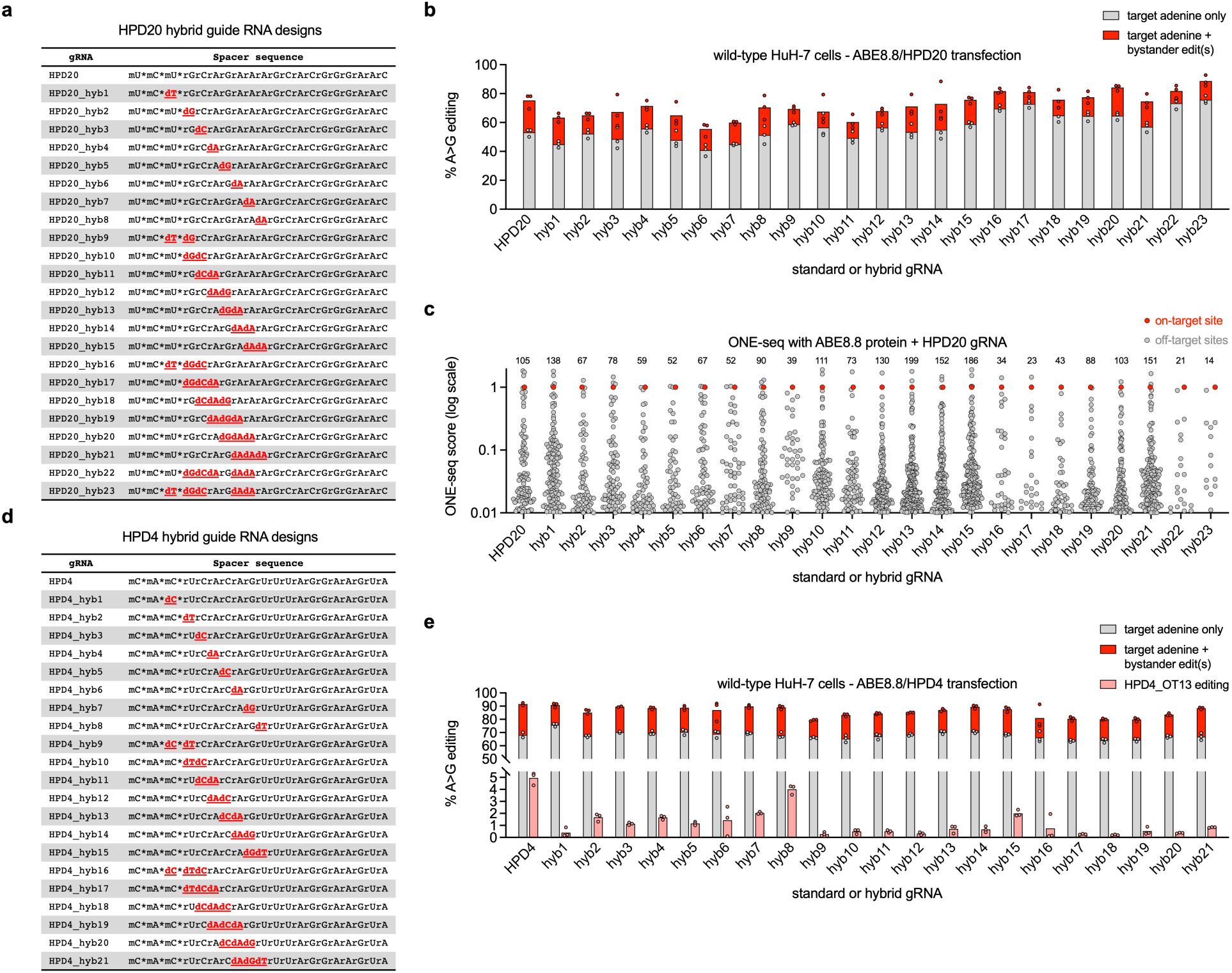
Optimizing targeting of two distinct sites in the *HPD* gene with hybrid gRNAs. **a**, Spacer sequences of HPD20 standard and hybrid gRNAs. “rN” = ribonucleotide; “dN” = deoxyribonucleotide; “m” = 2’-O-methylation; * = phosphorothioate linkage. DNA substitutions are in red bold and underlined. **b**, A-to-G editing of *HPD* on-target site (target adenine only, or target adenine plus bystander editing) in wild-type HuH-7 cells treated with HPD20 standard or hybrid gRNA in combination with ABE8.8 mRNA (*n* = 3 biological replicates per condition). **c**, Number of sites with ONE-seq score > 0.01 for each indicated HPD20 standard or hybrid gRNA with ABE8.8 protein. **d**, Spacer sequences of HPD4 standard and hybrid gRNAs. **e**, A-to-G editing of *HPD* on-target site (target adenine only, or target adenine plus bystander editing) or verified HPD4 off-target site (HPD4_OT13) in wild-type HuH-7 cells treated with HPD4 standard or hybrid gRNA in combination with ABE8.8 mRNA (*n* = 3 biological replicates per condition). Means are shown.

Because of its more favorable on-target and off-target editing profiles, we prioritized HPD20 for further study as a clinical lead gRNA. We assessed a full series of HPD20 hybrid gRNAs (HPD20_hyb1 through HPD20_hyb21) (**Fig. 4a**), matching the designs of the PAH1 hybrid gRNAs, via ABE8.8 mRNA/gRNA transfection in wild-type HuH-7 hepatocytes, for both editing of the target splice site adenine—in position 6 of the protospacer sequence, TCTGC**<u>A</u>**GAAAGCACGGGAAC, with a GGG PAM—and bystander editing of positions 8, 9, and/or 10 (**Fig. 4b**). Because we had not verified any individual genomic sites of off-target editing with the HPD20 standard gRNA, we used ONE-seq to assess the off-target editing profiles of the HPD20 hybrid gRNAs, with many showing decreased off-target potential (i.e., number of sites with ONE-seq scores greater than 0.01) but a few showing increased off-target potential (**Fig. 4c**). Two hybrid gRNAs, HPD20_hyb22 and HPD20_hyb23, with 5 DNA nucleotide substitutions each, maintained on-target editing, substantially reduced bystander editing, and had the fewest sites with ONE-seq scores greater than 0.01 (**Fig. 4a–c**).

We also assessed the HPD4 gRNA, having the protospacer sequence CACTC**<u>A</u>**CAGTTTAGGAAGTA and a GGG PAM, with the target adenine base in position 6 and a bystander adenine base in position 8. Having verified a site of off-target editing with the HPD4 standard gRNA (HPD4_OT13) with mean 4.9% editing, we assessed a full series of HPD4 hybrid gRNAs (HPD4_hyb1 through HPD4_hyb21) (**Fig. 4d**), matching the designs of the PAH1 hybrid gRNAs, via ABE8.8 mRNA/gRNA transfection in wild-type HuH-7 hepatocytes (**Fig. 4e**). All the hybrid gRNAs reduced OT13 off-target editing, some close to the background level, and many hybrid gRNAs reduced bystander editing while maintaining on-target editing.

### Optimizing correction of the *PAH* R408W variant with a PAM-relaxed ABE

In a previous study, we characterized a gRNA (herein termed “PAH4”) that corrects the *PAH* R408W variant *in cellulo* and *in vivo*^5^. Because there are no NGG PAMs situated appropriately for adenine base editing of the *PAH* R408W variant, we screened PAM-altered and PAM-relaxed versions^22^ of ABEs and identified an optimal combination of SpRY-ABE8.8 and the protospacer sequence GGCC**<u>A</u>**AGGTATTGTGGCAGC with an AAA PAM (**Fig. 1a**). The variant adenine base lies in position 5 of the protospacer sequence. Potential bystander adenine bases lie in positions 6 and 10; bystander editing at position 6 would result in a benign synonymous variant, whereas editing at position 10 would result in a nonsynonymous variant that has been reported to be a pathogenic variant for PKU^23,24^.

Editors based on the SpRY variant of *Streptococcus pyogenes* Cas9 are near-PAMless in their ability to engage genomic sites^22^, greatly expanding their targeting range but also incurring an increased potential for off-target editing. To evaluate the SpRY-ABE8.8/PAH4 combination, which corrects the *PAH* R408W variant, we performed ONE-seq against the reference human genome. Rather than interrogate candidate off-target sites nominated by ONE-seq with the PAH4 standard gRNA, we immediately tested a limited series of hybrid gRNAs (double/triple substitutions, PAH4_hyb9 through PAH4_hyb21, matching the designs of the analogous PAH1 hybrid gRNAs) with the goal of reducing the potential of SpRY-ABE8.8 for off-target editing and prospectively winnowing the list of candidate off-target sites for verification (**Fig. 5a**).

**Fig. 5.**
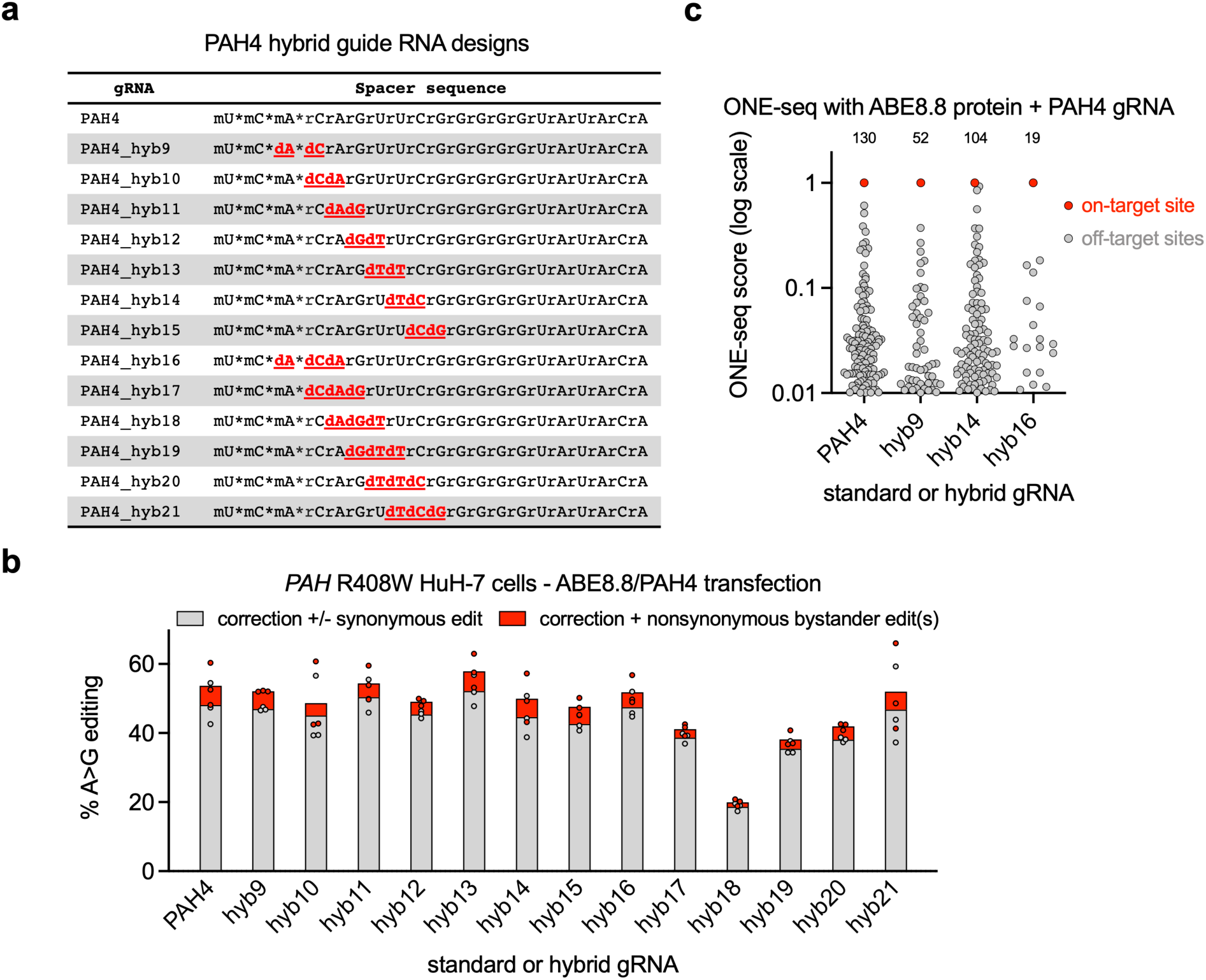
Optimizing correction of the *PAH* R408W variant with a PAM-relaxed ABE. **a**, Spacer sequences of PAH4 standard and hybrid gRNAs. “rN” = ribonucleotide; “dN” = deoxyribonucleotide; “m” = 2’-O-methylation; * = phosphorothioate linkage. DNA substitutions are in red bold and underlined. **b**, A-to-G editing of *PAH* on-target site (correction of R408W variant with or without synonymous bystander editing, or correction plus unwanted nonsynonymous bystander editing) in *PAH* R408W HuH-7 cells treated with PAH4 standard or hybrid gRNA in combination with SpRY-ABE8.8 mRNA (*n* = 3 biological replicates per condition). **c**, Number of sites with ONE-seq score > 0.01 for each indicated PAH4 standard or hybrid gRNA with SpRY-ABE8.8 protein.

Several of the PAH4 hybrid gRNAs displayed increased on-target editing and/or decreased bystander editing at protospacer position 10 (nonsynonymous variant) (**Fig. 5b**); ONE-seq demonstrated reduced off-target potential with some of these favorable hybrid gRNAs despite the use of a PAM-relaxed ABE (**Fig. 5c**).

## Discussion

In this study, we investigated the off-target editing profiles of clinical lead gRNAs in combination with ABEs for the correction of pathogenic variants in PKU and PXE and for inactivation of the *HPD* gene, a modifier for HT1. We assessed the ability of hybrid gRNAs to mitigate the potential for off-target editing by the standard versions of the gRNAs and found that some hybrid gRNAs can dramatically reduce off-target editing while also reducing bystander editing while maintaining or even enhancing on-target editing efficiency *in cellulo* and *in vivo*. Our findings suggest that the use of hybrid gRNAs to reduce off-target and bystander editing by ABEs is generalizable across different target loci and underscores its potential as a broadly applicable strategy to improve the safety and efficiency of base-editing therapies for various genetic disorders.

We note the limitations of this study. All the work in this study focused on adenine base editing; it is possible that the advantages we observed with the use of hybrid gRNAs *in cellulo* and *in vivo* are relevant to other editing modalities that rely on the use of gRNAs, such as nuclease editing, prime editing, and epigenome editing, but this would require validation for each modality. We present data for five distinct loci *in cellulo* and two of the loci *in vivo*, and in each case we were able to identify hybrid gRNAs that had favorable effects on editing specificity and efficiency, but it is possible that hybrid gRNAs would not be beneficial at all loci. In comparing our results across the five loci, our findings suggest that there will not be a single set of hybrid gRNA modifications that will fully optimize off-target and on-target editing across all genomic loci, although suggestive patterns are evident—e.g., the “hyb16” design (DNA nucleotide substitutions in protospacer positions 3, 4, and 5) substantially reduced off-target editing potential while maintaining on-target editing at all five tested loci, and so this design might be a reasonable starting point for future hybrid gRNA screening campaigns. Nonetheless, empirical testing will be needed for each new lead gRNA being considered for therapeutic use, at least for the foreseeable future. Although we have demonstrated that the use of a hybrid gRNA can substantially reduce gRNA-dependent off-target editing by an ABE both *in cellulo* and *in vivo*, it is not clear that use of a hybrid gRNA would affect gRNA-independent off-target editing, i.e., spurious RNA or DNA deamination by the ABE’s adenosine deaminase domain^25,26^; further experimentation would be needed to evaluate this possibility. These issues notwithstanding, it is clear that hybrid gRNAs can substantially improve some mRNA-LNP adenine base editing therapies—such as the ABE8.8/PAH1_hyb24 combination for the treatment of PKU, which we intend to take forward into an early-phase clinical trial in the near future.

## Methods

The research described here complied with all relevant regulations. All recombinant DNA research was approved by the Institutional Biosafety Committee at the University of Pennsylvania, where the studies were performed, and were consistent with local, state, and federal regulations as applicable, including the National Institutes of Health Guidelines for Research Involving Recombinant of Synthetic Nucleic Acid Molecules. All procedures used in animal studies were approved by the Institutional Animal Care and Use Committee at the University of Pennsylvania (protocol #805887), where the studies were performed, and were consistent with local, state, and federal regulations as applicable, including the National Institutes of Health Guide for the Care and Use of Laboratory Animals.

### RNA production

100-mer gRNAs were chemically synthesized under solid phase synthesis conditions by a commercial supplier (Agilent or Integrated DNA Technologies) with end-modifications as well as heavy 2’-O-methylribosugar modification as previously described^27^: for cellular studies, 5’-mX*mX*mX*XXXXXXXXXXXXXXXXXGUUUUAGAGCUAGAAAUAGCAAGUUAAA AUAAGGCUAGUCCGUUAUCAACUUGAAAAAGUGGCACCGAGUCGGUGCU*U*U*U-3’; for mouse studies, 5’-mX*mX*mX*XXXXXXXXXXXXXXXXXGUUUUAGAmGmCmUmAmGmAmAmAmUmA mGmCAAGUUAAAAUAAGGCUAGUCCGUUAUCAmAmCmUmUmGmAmAmAmAmAmGmUmGmGmCmAmCmCmGmAmGmUmCmGmGmUmGmCmU*mU*mU*mU-3’; where the Xs indicate the spacer sequence positions (specified in Fig. 2b, Fig. 3a, Fig. 4a, Fig. 4d, and Fig. 5a), and “m” and * respectively indicate 2’-O-methylation and phosphorothioate linkage.

ABE8.8 mRNA and SpRY-ABE8.8 mRNA were produced via in vitro transcription (IVT) and purification as previously described^4,5^. In brief, a plasmid DNA template containing a codon-optimized ABE8.8 or SpRY-ABE8.8 coding sequence and a 3’ polyadenylate sequence was linearized. An IVT reaction containing linearized DNA template, T7 RNA polymerase, NTPs, and cap analog was performed to produce mRNA containing N1-methylpseudouridine. After digestion of the DNA template with DNase I, the mRNA product underwent purification and buffer exchange, and the purity of the final mRNA product was assessed with spectrophotometry and capillary gel electrophoresis. Elimination of double-stranded RNA contaminants was assessed using dot blots and transfection into human dendritic cells. Endotoxin content was measured using a chromogenic Limulus amebocyte lysate (LAL) assay; all assays were negative.

### LNP formulation

LNPs were formulated as previously described^4,5,28^, with the lipid components (SM-102, 1,2-distearoyl-sn-glycero-3-phosphocholine, cholesterol, and PEG-2000 at molar ratios of 50:10:38.5:1.5) being rapidly mixed with an aqueous buffer solution containing ABE8.8 mRNA and a gRNA in a 1:1 ratio by weight in 25 mM sodium acetate (pH 4.0), with an N:P ratio of 5.6. The resulting LNP formulations were subsequently dialyzed against sucrose-containing buffer, concentrated using Amicon Ultra-15 mL Centrifugal Filter Units (Millipore Sigma), sterile-filtered using 0.2-µm filters, and frozen until use. The LNPs underwent quality control for particle size (Z-Ave, hydrodynamic diameter), polydispersity index as determined by dynamic light scattering (Malvern NanoZS Zetasizer) and total RNA encapsulation as measured by the Quant-iT Ribogreen Assay (Thermo Fisher Scientific).

### Culture and transfection of HuH-7 cells

Wild-type HuH-7 cells, *PAH* P281L homozygous HuH-7 cells^4^, *PAH* R408W homozygous HuH-7 cells^5^, and *ABCC6* R1164X homozygous HuH-7 cells (**Supplementary Note 2**) were used for this study. HuH-7 cells were maintained in Dulbecco’s modified Eagle’s medium (containing 4 mM L-glutamine and 1 g/L glucose) with 10% fetal bovine serum and 1% penicillin/streptomycin at 37°C with 5% CO_2_. HuH-7 cells were seeded on 24-well plates (Corning) at 1.2 × 10^5^ cells per well. For plasmid transfections, at 3–4 hours after seeding, cells were transfected at approximately 80–90% confluency with 1.5 μL TransIT®-LT1 Transfection Reagent (MIR2300, Mirus), 0.25 μg base editor plasmid, and 0.25 μg gRNA plasmid per well according to the manufacturer’s instructions. An ABE8.8 (ABE8.8-m) expression plasmid^10^ was used as previously described^4^, and HPD3, HPD4, HPD5, HPD20, and PXE1 gRNA expression plasmids were newly generated for this study. For mRNA/gRNA transfections, in one tube, 3 µL Lipofectamine MessengerMax (LMRNA008, Thermo Fisher Scientific) was added to 50 µL of Opti-MEM and incubated for 10 minutes; in another tube, 0.5 µg mRNA and 0.5 µg gRNA was suspended in a total of 50 µL of Opti-MEM. The diluted RNA was added to the tube of diluted MessengerMAX, and the subsequent RNA/MessengerMax/Opti-MEM mixture was incubated for 5 minutes at room temperature and then added dropwise onto the cells. For each type of transfection, cells were cultured for 72 hours after transfection, and then media were removed, cells were washed with 1× DPBS (Corning), and genomic DNA was isolated using the DNeasy Blood and Tissue Kit (QIAGEN) according to the manufacturer’s instructions.

### ONE-seq

ONE-seq was performed as previously described^4,8,19^. The human ONE-seq libraries for the PAH1, PAH4, PXE1, HPD3/HPD4/HPD5 (combined library), and HPD20 gRNAs were designed using the GRCh38 Ensembl v98 reference genome (ftp://ftp.ensembl.org/pub/release-98/fasta/homo_sapiens/dna/Homo_sapiens.GRCh38.dna.chromosome.{1-22,X,Y,MT}.fa, ftp://ftp.ensembl.org/pub/release-98/fasta/homo_sapiens/dna/Homo_sapiens.GRCh38.dna.nonchromosomal.fa), and the mouse ONE-seq libraries for the PAH1 and PXE1 gRNAs were designed using the GRCm39 Ensembl reference genome (ftp://ftp.ensembl.org/pub/release-113/fasta/mus_musculus/dna/Mus_musculus.GRCm39.dna.chromosome.{1-19,X,Y,MT}.fa, ftp://ftp.ensembl.org/pub/release-113/fasta/mus_musculus/dna/Mus_musculus.GRCm39.dna.nonchromosomal.fa). Sites with up to 6 mismatches and sites with up to 4 mismatches plus up to 2 DNA or RNA bulges, compared to the on-target site, were identified with Cas-Designer v1.2^20^. The variant-aware ONE-seq library for the PAH1 gRNA was designed using CRISPRme^21^, with similar mismatch criteria. The final oligonucleotide sequences were generated with a script^19^, and the oligonucleotide libraries were synthesized by Twist Biosciences. Recombinant ABE8.8 protein and SpRY-ABE8.8 protein were produced by GenScript. Duplicate ONE-seq experiments were previously performed for ABE8.8/PAH1^4^; mean ONE-seq scores were used for this study. All other ONE-seq experiments were performed specifically for this study as follows. Each library was PCR-amplified and subjected to 1.25× AMPure XP bead purification. After incubation at 25°C for 10 minutes in CutSmart buffer, RNP comprising 769 nM recombinant ABE8.8-m protein and 1.54 µM gRNA was mixed with 100 ng of the purified library and incubated at 37°C for 8 hours.

Proteinase K was added to quench the reaction at 37°C for 45 minutes, followed by 2× AMPure XP bead purification. The reaction was then serially incubated with EndoV at 37°C for 30 minutes, Klenow Fragment (New England Biolabs) at 37°C for 30 minutes, and NEBNext Ultra II End Prep Enzyme Mix (New England Biolabs) at 20°C for 30 minutes followed by 65°C for 30 minutes, with 2× AMPure XP bead purification after each incubation. The reaction was ligated with an annealed adaptor oligonucleotide duplex at 20°C for 1 hour to facilitate PCR amplification of the cleaved library products, followed by 2× AMPure XP bead purification. Size selection of the ligated reaction was performed on a BluePippin system (Sage Science) to isolate DNA of 150-200 bp on a 3% agarose gel cassette, followed by two rounds of PCR amplification to generate a barcoded library, which underwent paired-end sequencing on an Illumina MiSeq System as described below. The analysis pipeline^19^ used for processing the data assigned a score quantifying the editing efficiency with respect to the on-target site to each potential off-target site. Sites were ranked based on this ONE-seq score, which was used for site prioritization.

### Next-generation sequencing (NGS)

NGS was performed as previously described^4^. For targeted amplicon sequencing, PCR reactions were performed using NEBNext Polymerase (NEB) using the primer sets listed in Supplementary Table 10, designed with Primer3 v4.1.0 (https://primer3.ut.ee/). The following program was used for all genomic DNA PCRs: 98°C for 20 seconds, 35× (98°C for 20 seconds, 57°C for 30 seconds, 72°C for 10 seconds), 72°C for 2 minutes. PCR products were visualized via capillary electrophoresis (QIAxcel, QIAGEN) and then purified and normalized via an NGS Normalization 96-Well Kit (Norgen Biotek Corporation). A secondary barcoding PCR was conducted to add Illumina barcodes (Nextera XT Index Kit V2 Set A and/or Nextera XT Index Kit V2 Set D) using ≈15 ng of first-round PCR product as template, followed by purification and normalization. Final pooled libraries were quantified using a Qubit 3.0 Fluorometer (Thermo Fisher Scientific) and then after denaturation, dilution to 10 pM, and supplementation with 15% PhiX, underwent single-end or paired-end sequencing on an Illumina MiSeq System. The amplicon sequencing data were analyzed with CRISPResso2 v2^29^ and scripts to quantify editing. For on-target editing, A-to-G editing was quantified at the site of the target adenine and at the site(s) of each potential bystander adenine in the editing window; for candidate off-target sites, A-to-G editing was quantified throughout the editing window (positions 1 to 10 of the protospacer sequence). In some cases, PCR amplicons were subjected to confirmatory Sanger sequencing, performed by GENEWIZ.

### Hybrid capture sequencing

The PAH1 probe library was generated to cover the 280 ONE-seq-nominated genomic sites with ONE-seq scores greater than 0.01, as well as additional sites. The probe library was generated serially using Agilent SureDesign, where probes were created using three rounds of generation settings. In the initial round, probes were generated with 2× tiling density, moderately stringent masking, and optimized performance with 90 minutes hybridization boosting. Regions that were not covered with the initial settings had probes generated with 2× tiling density, least stringent masking, and optimized performance with 90 minutes hybridization boosting. The remaining uncovered regions had probes generated with 1× tiling density, no masking, and no boosting. The hybrid capture procedure was conducted using the SureSelect XT HS2 DNA Reagent Kit (Agilent) as per the manufacturer’s instructions.

Briefly, 200 ng of genomic DNA from ABE8.8/PAH1 mRNA/gRNA-treated or control untreated HuH-7 cells, quantified on the TapeStation Genomic DNA ScreenTape platform (Agilent), were prepared and enzymatically fragmented with modifications for 2×150 reads. The DNA fragments were then processed for library construction using molecular barcodes. Adaptors were ligated to the DNA fragments, followed by repair, dA-tailing, and purification using AMPure XP beads. The libraries were amplified (8 cycles), indexed, and purified with AMPure XP beads, and their quality was assessed via the TapeStation D1000 ScreenTape platform (Agilent); 1000 ng of each library was prepared and hybridized to the PAH1 probe library, captured using streptavidin beads, amplified (16 cycles), and purified using AMPure XP beads. The final libraries were purified and quantified via the TapeStation High Sensitivity D1000 ScreenTape platform (Agilent). The libraries were pooled and sequenced on an Illumina NovaSeq, performed by Novogene. The resulting reads were processed and analyzed using the Agilent Alissa Reporter software with the following settings: normal/tumor pair analysis mode for treated samples, and tumor only for control samples, molecular barcode deduplication mode, duplex MBC deduplication consensus mode, 2 read pairs per MBC minimum, 10 reads minimum coverage depth, 3 reads minimum supporting variant allele, and 0.001 minimum variant allele frequency. For each of the 280 sites, A-to-G editing was quantified throughout the editing window (Supplementary Table 1). For sites with any position displaying a mean net A-to-G editing level ≥ 0.1% in treated versus control samples, the sites were confirmed with targeted amplicon sequencing, as described above. Only 2 out of the 280 sites failed to be captured and sequenced, with both sites being on chromosome Y, which is absent from HuH-7 cells.

### Mouse studies

Mouse studies were performed as previously described^4^. Mice were maintained on a 12-hour light/12-hour dark cycle, with a temperature range of 65°F to 75°F and a humidity range of 40% to 60%, and were fed ad libitum with a chow diet (LabDiet, Laboratory Autoclavable Rodent Diet 5010). Humanized PKU mice homozygous for the *PAH* P281L allele^4^ or for the *ABCC6* R1164X allele (Supplementary Note 3) on the C57BL/6J background were generated as littermates/colonymates with wild-type C57BL/6J mice. Genotyping was performed using PCR amplification from genomic DNA samples (prepared from clipped tails/ears) followed by Sanger sequencing or NGS. Roughly equal numbers of female and male littermates/colonymates were used for experiments at ages ranging from six to eight weeks, with random assignment of animals to various experimental groups when applicable, and with collection and analysis of data performed in a blinded fashion when possible. LNPs were administered to mice at a dose of 2.5 mg/kg via retro-orbital injection under anesthesia with 1%-2% inhaled isoflurane. Blood samples were collected via the tail tip at various timepoints: pre-treatment or 4 hours, 24 hours, 48 hours, or 7 days after treatment. Mice were euthanized 1 to 2 weeks after treatment, and 8 liver samples (2 from each lobe) were obtained on necropsy and processed with the DNeasy Blood and Tissue Kit (QIAGEN) as per the manufacturer’s instructions to isolate genomic DNA. Euthanasia in all instances was achieved via terminal inhalation of carbon dioxide followed by secondary euthanasia through cervical dislocation or decapitation, consistent with the 2020 American Veterinary Medical Association Guidelines on Euthanasia. NGS results from the liver samples were averaged to provide quantification of whole-liver editing.

### Measurement of blood phenylalanine

Blood phenylalanine levels were measured by an enzymatic method using the Phenylalanine Assay Kit (MAK005, Millipore Sigma) according to the manufacturers’ instructions. Briefly, plasma samples were collected at timepoints in the early afternoon, to account for diurnal variation in blood phenylalanine levels, and deproteinized with a 10 kDa MWCO spin filter (CLS431478-25EA, Millipore Sigma) and pre-treated with 5 µL of tyrosinase for 10 minutes at room temperature prior to start of the assay. Reaction mixes were made according to the manufacturers’ instructions, and the fluorescence intensity of each sample was measured (λ_ex_ = 535/λ_em_ = 587 nm).

### Measurement of blood pyrophosphate

Plasma samples collected pre-treatment and 7 days post-treatment were diluted 1:1 in Tris-acetate pH 8.0 and centrifuged with a 30K filter (OD030C34, Cytiva) at 14,000 × g at 4°C for 20 minutes. The pyrophosphate concentrations were measured in platelet-free plasma using ATP sulfurylase to convert pyrophosphate into ATP in the presence of excess adenosine 5’ phosphosulfate followed by a luminescent assay of ATP, as described previously^30,31^.

### Measurement of ALT, cytokines, and chemokines

Assays were performed as previously described^5^. Alanine aminotransferase (ALT) (MAK052-1KT, Millipore Sigma) activity was measured according to the manufacturer’s instructions. The expression profile of cytokines and chemokines in mouse serum samples was determined using the BioLegend anti-virus response panel [13-plex bead-based assay for the quantification of interferons (α, β, γ), interleukins (1β, 6, 10, 12p70), and chemokines (MCP-1, RANTES, CXCL-1, IP-10, TNF-α, and GM-CSF)] according to the manufacturer’s instructions. Briefly, kit components were thawed and reconstituted as needed. Standards were prepared by 1:4 serial dilution in assay buffer. Serum samples were diluted 1:3 in Matrix A. Filter bottom plates were prewetted with wash buffer before addition of standards and samples. 25 µL of each standard and sample was added to the plates in duplicates and mixed with 25 µL of premixed beads solution. Plates were incubated with shaking at 500 rpm for 2 hours and washed twice, followed by addition of 25 µL detection antibodies and incubation at 500 rpm for 1 hour. Without washing, 25 µL SA-PE was added to each well and incubated with shaking at 500 rpm for 30 minutes. Plates were washed twice, and beads were resuspended in 150 µL wash buffer, transferred to FACS tubes, and acquired on an BD LSR flow cytometer. The standard curve was used to determine the concentrations of each analyte in the serum samples using a 5PL fitting.

## Data analysis

Sequencing data were analyzed as described above. Other data were collected and analyzed using GraphPad Prism v10.4.1.

## Data availability

DNA sequencing data that support the findings of this study will be deposited in the NCBI Sequence Read Archive upon publication of this manuscript. Other data supporting the findings of this study are available within the manuscript and its Supplementary Information. The GRCh38 Ensembl v98 reference genome (ftp://ftp.ensembl.org/pub/release-98/fasta/homo_sapiens/dna/Homo_sapiens.GRCh38.dna.chromosome.{1-22,X,Y,MT}.fa, ftp://ftp.ensembl.org/pub/release-98/fasta/homo_sapiens/dna/Homo_sapiens.GRCh38.dna.nonchromosomal.fa) annotation and the GRCm39 Ensembl reference genome (ftp://ftp.ensembl.org/pub/release-113/fasta/mus_musculus/dna/Mus_musculus.GRCm39.dna.chromosome.{1-19,X,Y,MT}.fa, ftp://ftp.ensembl.org/pub/release-113/fasta/mus_musculus/dna/Mus_musculus.GRCm39.dna.nonchromosomal.fa) annotation were used.

## Acknowledgements

This work was supported by U.S. National Institutes of Health (NIH) grant U19-NS132301 (W.H.P., R.C.A.-N., K.M., M.-G.A., and X.W.) through the Common Fund Program, Somatic Cell Genome Editing, by NIH grant R35-HL145203 (K.M.), by NIH grant R01-HD110733 and an American Heart Association Career Development Award (X.W.), by NIH grant R21-AR084212 (Q.L.), by a Graduate Research Fellowship from the U.S. National Science Foundation and an American Heart Association Predoctoral Fellowship (M.N.W.), and by the Winkelman Family Fund in Cardiovascular Innovation (K.M.). The authors wish to acknowledge assistance from Jason Fischer, Taylor Jones, and Briana Kipphut at Agilent Technologies, with design of probe libraries and data analysis related to hybrid capture sequencing, and Archana Lovett, Uma Nagarajan, and Barrie Rogers at Novogene Corporation, with sequencing of hybrid capture libraries.

## Author contributions

K.M., M.-G.A., W.H.P., and X.W. supervised the work with intellectual input on experimental design and data analysis from J.Z.W., R.C.A.-N., and Q.L. M.N.W., L.C.T., D.L.B., A.Q., I.J., D.V., P.Q., Q.L., and X.W. contributed to web laboratory experiments. M.N.W. and K.M. contributed to bioinformatic analyses. H.S, G.D., and M.-G.A. contributed to mRNA production, LNP formulation, and cytokine/chemokine analysis. M.N.W., L.C.T., K.M., and X.W. drafted the manuscript, and all authors contributed to the editing of the manuscript.

---

**Extended Fig. 1.**
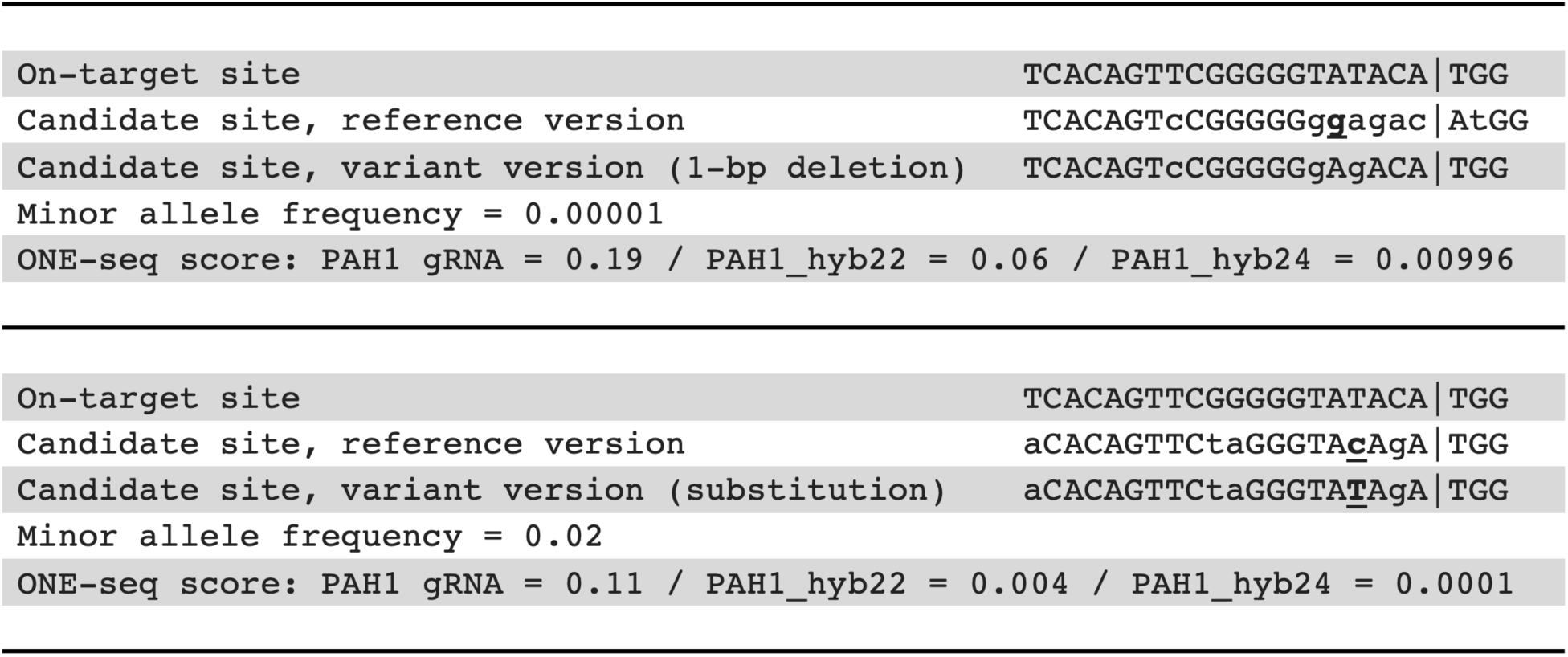
Reduction of variant-aware off-target editing by PAH1 hybrid guide RNAs. Examples of variant sites nominated by ONE-seq, with reductions in ONE-seq scores observed with PAH1 hybrid gRNAs. Differences between reference and variant versions of nominated sites are in bold and underlined.

**Extended Fig. 2.**
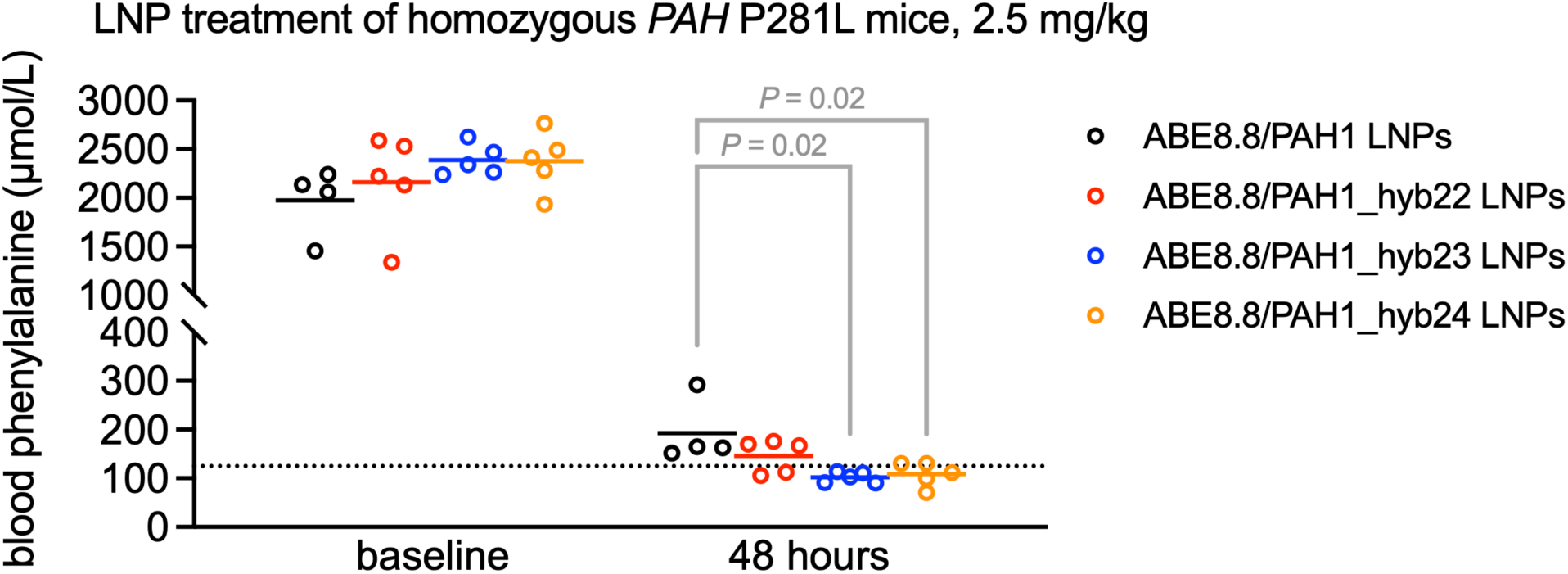
Reduction of blood phenylalanine levels by PAH1 hybrid guide RNAs. Blood phenylalanine levels in homozygous *PAH* P281L mice (*n* = 4–5 animals per group) before and 48 hours after treatment with 2.5 mg/kg dose of LNPs with PAH1 standard or hybrid gRNA in combination with ABE8.8 mRNA (1 blood sample per timepoint). The dotted line marks the upper end of the normal range for blood phenylalanine, 125 µmol/L. Means are shown; *P* values were calculated with the Mann–Whitney *U* test.

**Extended Fig. 3.**
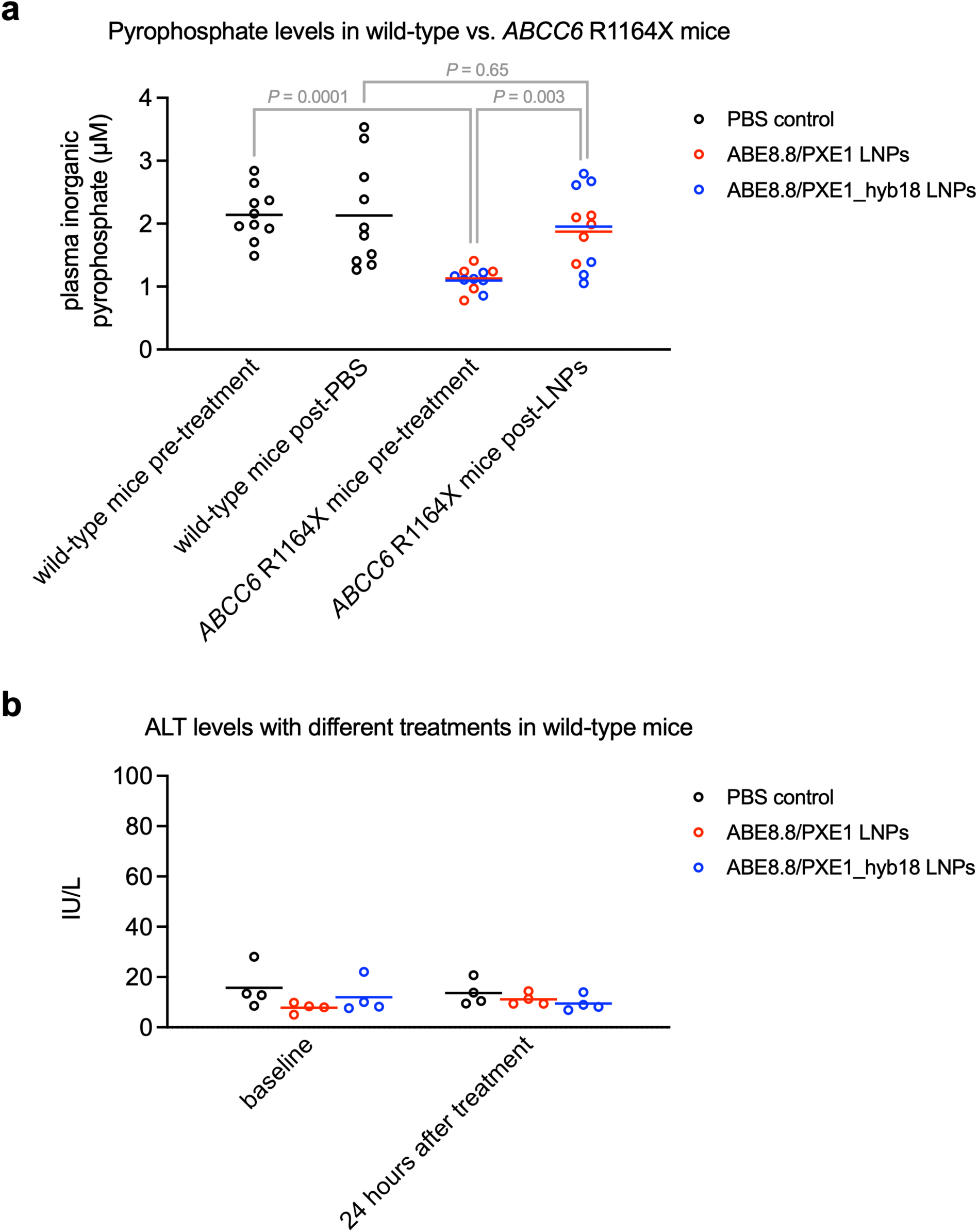
Effects of PXE1 gRNA LNPs on blood pyrophosphate levels and ALT levels. **a**, Blood pyrophosphate levels in wild-type mice, before and 7 days after treatment with PBS vehicle (*n* = 10 animals), or in homozygous *ABCC6* R1164X mice, before and 7 days after treatment with 2.5 mg/kg dose of LNPs with PAH1 standard gRNA (*n* = 6 animals) or PAH1_hyb18 gRNA (*n* = 5 animals) in combination with ABE8.8 mRNA (1 blood sample per timepoint). Means are shown; *P* values comparing genotype groups were calculated with the Mann–Whitney *U* test, and the *P* value comparison for the aggregated R1164X groups (before versus after treatment) was calculated with the Wilcoxon signed-rank test. *P* values were not calculated within each individual R1164X gRNA group (before versus after treatment) because of inadequate sample sizes for the Wilcoxon signed-rank test. **b**, Blood alanine aminotransferase (ALT) levels in wild-type mice (*n* = 4 animals per group) before and 24 hours after treatment with PBS vehicle or with 2.5 mg/kg dose of LNPs with PAH1 standard or hybrid gRNA in combination with ABE8.8 mRNA (1 blood sample per timepoint). No effects on ALT levels by LNP treatment were evident.

**Extended Figure 4.**
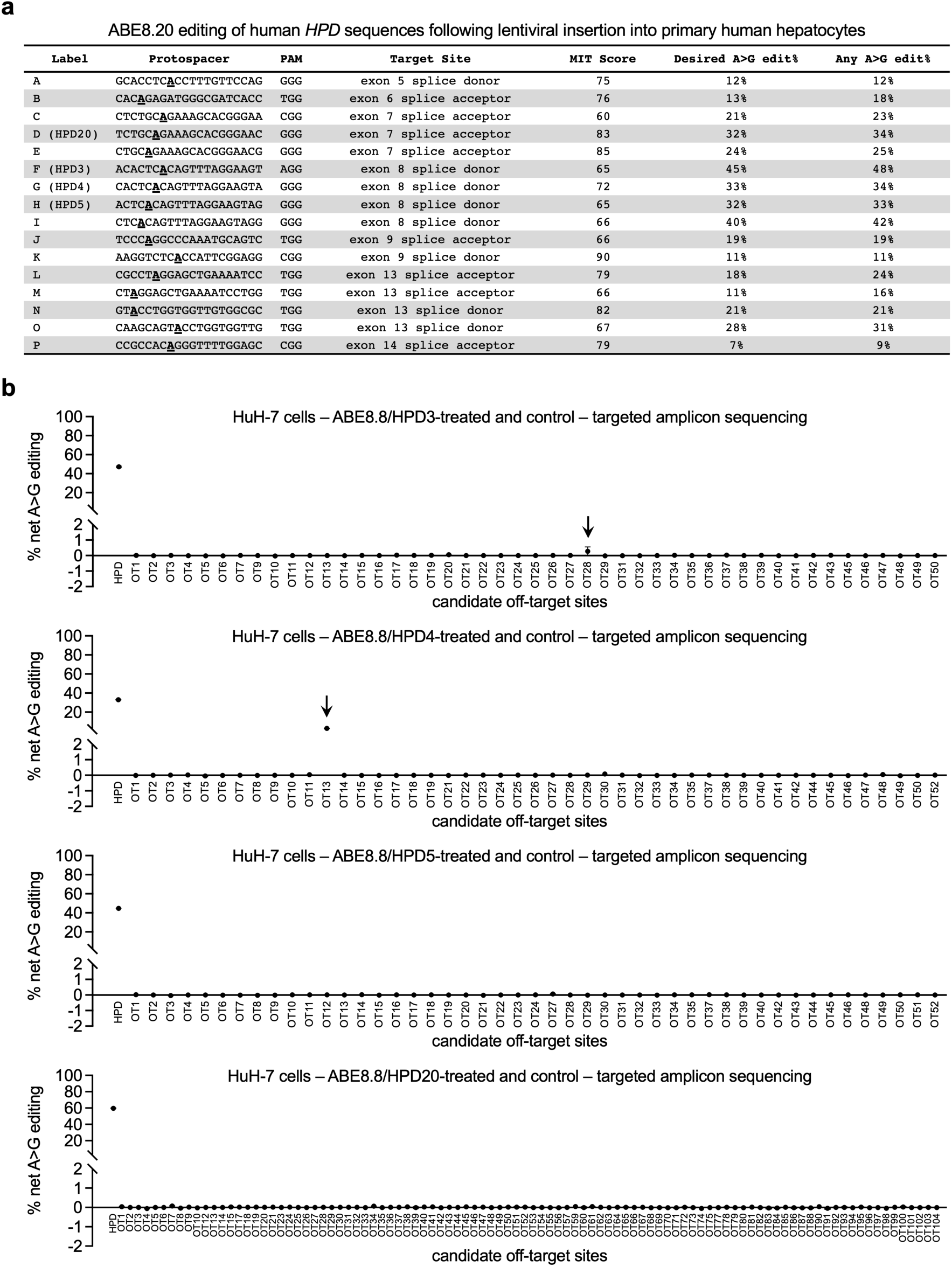
Assessment of on-target and off-target editing by guide RNAs targeting the *HPD* gene. **a**, Lentiviral screening in primary human hepatocytes for adenine base editing of heterologous *HPD* splice site target sequences by various gRNAs in combination with ABE8.20; target adenines are in bold and underlined. MIT score = off-target/specificity score indicated by UCSC Genome Browser. Four candidate gRNAs (HPD3, HPD4, HPD5, and HPD20) were taken forward for off-target analyses. **b**, Targeted amplicon sequencing of ONE-seq-nominated sites in plasmid-treated vs. control wild-type HuH-7 cells with HPD3, HPD4, HPD5, or HPD20 gRNA in combination with ABE8.8 (*n* = 2 treated and 2 untreated biological replicates). Sites with unsuccessful sequencing are omitted. Arrows indicate verified sites of off-target editing. Means ± standard deviations are shown.

## Supplementary Note 1

For the PAH1 gRNA, none of the seven verified sites of off-target editing raises a concern of oncogenic risk. The first site (PAH1_OT1) has 3 mismatches with the on-target protospacer, with an NGG PAM, and is in an intron of the *SPAG17* gene, which encodes a microtubule protein. PAH1_OT2 has 5 mismatches with the on-target protospacer, with an NGA PAM, and is in an intron of the *STYXL2* gene, which encodes a myosin-interacting protein. PAH1_OT3 has 2 mismatches and 2 consecutive bulges compared with the on-target protospacer, with an NGG PAM, and is in an intron of *LINC00882*, a long noncoding RNA. PAH1_OT4 has 6 mismatches with the on-target protospacer, with an NGG PAM, and is in an intron of DCLK1, which encodes a microtubule-associated protein. PAH1_OT5 and PAH1_OT6 each have 4 mismatches with the on-target protospacer, with an NGG PAM, and is intergenic and distant from any gene. PAH1_OT7 has 4 mismatches and 2 consecutive bulges compared with the on-target protospacer, with an NGG PAM, and is in an intron of *KCNJ10*, which encodes a potassium channel protein. None of the five implicated genes appear to be significantly expressed in the liver.

For the PXE1 gRNA, none of the six verified sites of off-target editing raises a concern of oncogenic risk. The first site (PXE1_OT1) has 2 mismatches and 1 bulge compared with the on-target protospacer, with an NGA PAM, and is intergenic and distant from any gene. PXE1_OT2 has 4 mismatches with the on-target protospacer, with an NGG PAM, and is in an intron of the *FGGY* gene, which encodes a carbohydrate kinase domain-containing protein expressed in fetal brain, with low expression in the liver. PXE1_OT8 has 4 mismatches with the on-target protospacer, with an NGG PAM, and is intergenic and distant from any gene. PXE1_OT10 has 3 mismatches with the on-target protospacer, with an NGG PAM, and is intergenic and distant from any gene. PXE1_OT14 has 6 mismatches with the on-target protospacer, with an NGG PAM, and is in the 5’ untranslated region of the *KCNK2* gene, which encodes a neuronal potassium channel protein. PXE1_OT15 has 6 mismatches with the on-target protospacer, with an NGG PAM, and is intergenic and distant from any gene.

For the HPD4 gRNA, the verified site of off-target editing raises no concern of oncogenic risk. HPD4_OT13 has 2 mismatches and 1 bulge compared with the on-target protospacer, with an NGG PAM, and is intergenic and distant from any gene.

## Supplementary Note 2

We used prime editing, specifically PE5max with an engineered pegRNA (epegRNA), to introduce the *ABCC6* R1164X variant into HuH-7 human hepatoma cells. HuH-7 cells in a well of a 6-well plate were transfected with 9 μL TransIT®-LT1 Transfection Reagent, 1.5 μg PEmax plasmid (pCMV-PEmax-P2A-hMLH1dn; Addgene plasmid # 174828; http://n2t.net/addgene:174828; RRID:Addgene_174828), 0.75 µg epegRNA-expressing plasmid (pU6-tevopreq1-GG-acceptor; Addgene plasmid # 174038; http://n2t.net/addgene:174038; RRID:Addgene_174038; spacer sequence 5’-ACCUGUCAGCCACCAG UCGC-3’; RTT/PBS sequence 5’-AGUUUCCCG**<u>U</u>**GACUGGUGGCUGA-3’; the bold underlined position introduces the *ABCC6* R1164X variant), and 0.75 μg nicking gRNA plasmid (pGuide; Addgene plasmid # 64711; http://n2t.net/addgene:64711; RRID:Addgene_64711; spacer sequence 5’-GAUCAGU UUCCCGUGACUGG-3’) as previously described^4^. Observing ≈30% editing in bulk HuH-7 cells, we used single-cell cloning to identify and expand a R1164X homozygous HuH-7 cell line. Cells were dissociated with trypsin 48 hours post-transfection and replated onto 10-cm plates (5,000 cells/plate) with conditioned medium to facilitate recovery, and genomic DNA was isolated from the remainder of the cells as a pool to perform PCR and Sanger sequencing of the *ABCC6* R1164X site. Single cells were permitted to expand for 7-14 days to establish clonal populations. Colonies were manually picked and replated into individual wells of a 96-well plate. Genomic DNA was isolated from individual clones, and PCR and Sanger sequencing was performed to identify R1164X homozygous HuH-7 clones. One representative clone was expanded for use in subsequent studies. This R1164X homozygous HuH-7 cell line is readily available from the authors via a Transfer of Research Material agreement with the University of Pennsylvania.

## Supplementary Note 3

The *ABCC6* R1164X mouse model of PXE was generated using in vitro transcribed Cas9 mRNA, a synthetic gRNA (spacer sequence 5’-GAGGGUCAGUUUCCCGAAAC-3’) (Integrated DNA Technologies), and a synthetic single-strand DNA oligonucleotide (Integrated DNA Technologies) with homology arms matching the target site and harboring the R1164X variant and synonymous variants (bold with underline): 5’-TCCAGGGAAGTCTGGTGGTCAGGGCCTTCCGGGCCCAGGCATCCTT CACGGCTCAGCACGATGCTCTCATGGATGAGAACCAGAGG**<u>A</u>**TCAGTTTCCCG**<u>TG</u>**ACTGGTG GCTGACAGGTAGGATGAGCCTGGGATCAGAGTAGATCTGGAACCCACTCCCCACAAGAGCTGCACTCTGTAGTTAGAGGGGTCACATGGCTTG-3’. The mixture of the 3 components was injected into cytoplasm of fertilized oocytes from C57BL/6J mice at the Penn Vet Transgenic Mouse Core (https://www.vet.upenn.edu/research/core-resources-facilities/transgenic-mouse-core). Genomic DNA samples from founders were screened for knock-in of the desired sequence in the *Abcc6* locus via homology-directed repair. Founders with the humanized R1164X allele were bred through two generations to obtain homozygous mice. This humanized PXE mouse model is readily available from the authors via a Transfer of Research Material agreement with the University of Pennsylvania.

## Supplementary Note 4

We implemented a high-throughput screening platform in which to introduce a base editor in combination with a variety of gRNAs and rapidly evaluate the relative activities of the gRNAs in the target cell type of interest, primary human hepatocytes. We used lentiviral vectors harboring an expression cassette for an adenine base editor and a combination of an expression cassette for an individual gRNA and a genomic fragment that (1) spanned the potential target sequence of that gRNA, (2) included a unique barcode, and (3) was flanked by fixed sequences that allowed for PCR amplification with a common set of primers. For the screen, we chose ABE8.20, being a high-activity eighth-generation adenine base editor with a wide editing window (protospacer positions 3 to 9)^10^ that permitted the greatest flexibility in gRNA selection for the disruption of splice donors (editing of the antisense A in the second position of the GT motif) or splice acceptors (editing of the sense A in the first position of the AG motif) in the *HPD* gene. We selected 16 gRNAs matched to protospacer sequences in the human *HPD* gene and with NGG PAMs, targeting splice sites ranged across the entire HPD gene (Extended Fig. 4a).

A pool of oligonucleotides spanning guide RNA sequences and target sequences, along with unique barcodes, was synthesized by Twist Bioscience and was amplified using KAPA HiFi HotStart ReadyMix (Roche), using various conditions to minimize template switching. After determining the optimal PCR strategy (48 parallel PCR reactions with 0.5 ng of the oligonucleotide pool per reaction, with 15 cycles), the amplified pool was cloned into a lentiviral vector using In-Fusion (Takara Bio) per the manufacturer’s instructions. Lentiviruses with a gene encoding ABE8.20 (ABE8.20-m^10^) and with the cloned gRNA/barcode/target sequences were generated using the Broad Institute’s RNAi Consortium protocol (https://portals.broadinstitute.org/gpp/public/dir/download?dirpath=protocols/ production&filename=TRC%20shRNA%20sgRNA%20ORF%20Low%20Throughput%20Viral%20Pro duction%20201506.pdf) for low-throughput lentiviral production and then concentrated using PEG-it virus precipitation solution (System Biosciences) per the manufacturer’s instructions. Primary human hepatocytes (Gibco/Thermo Fisher Scientific) were thawed into a 24-well Collagen I coated plate (Gibco/Thermo Fisher Scientific) and transduced with the lentiviruses 6 hours later. Each well of cells was treated via addition of 1 µL of concentrated virus, with 8 biological replicates. Four days after transduction, each group was pooled and then harvested using the DNeasy Blood and Tissue Kit (QIAGEN). NGS library preparation was performed with KAPA HiFi HotStart ReadyMix (Roche), with 35 PCR cycles using a primer pair spanning the barcode and the target sequence. Following a secondary indexing PCR round and sequencing on an Illumina MiSeq system, target sequences were binned via the unique barcodes, and A-to-G editing was quantified in the editing window of each target sequence.

We observed the highest levels of editing with gRNAs targeting either the splice acceptor of exon 7 or the splice donor of exon 8 (Extended Fig. 4a). We prioritized the gRNAs designated “HPD3”, “HPD4”, “HPD5”, and “HPD20” due to their efficient editing. Noting that these gRNAs in combination with ABE8.20 achieved editing of adenine bases towards the middle of the editing window (positions 4 through 7), for our subsequent studies we switched to ABE8.8 due to its narrow editing window. We retested the HPD3, HPD4, HPD5, and HPD20 gRNAs in combination with ABE8.8, now using plasmid transfection of ABE8.8 and gRNA in wild-type HuH-7 hepatocytes (Extended Fig. 4b). All demonstrated efficient editing of their target splice site adenine, with HPD20 performing the best. Of note, all gRNAs demonstrated bystander editing, but such editing is expected to be of little consequence with respect to the inactivation of the *HPD* gene.

---

**Supplementary Table 2.**
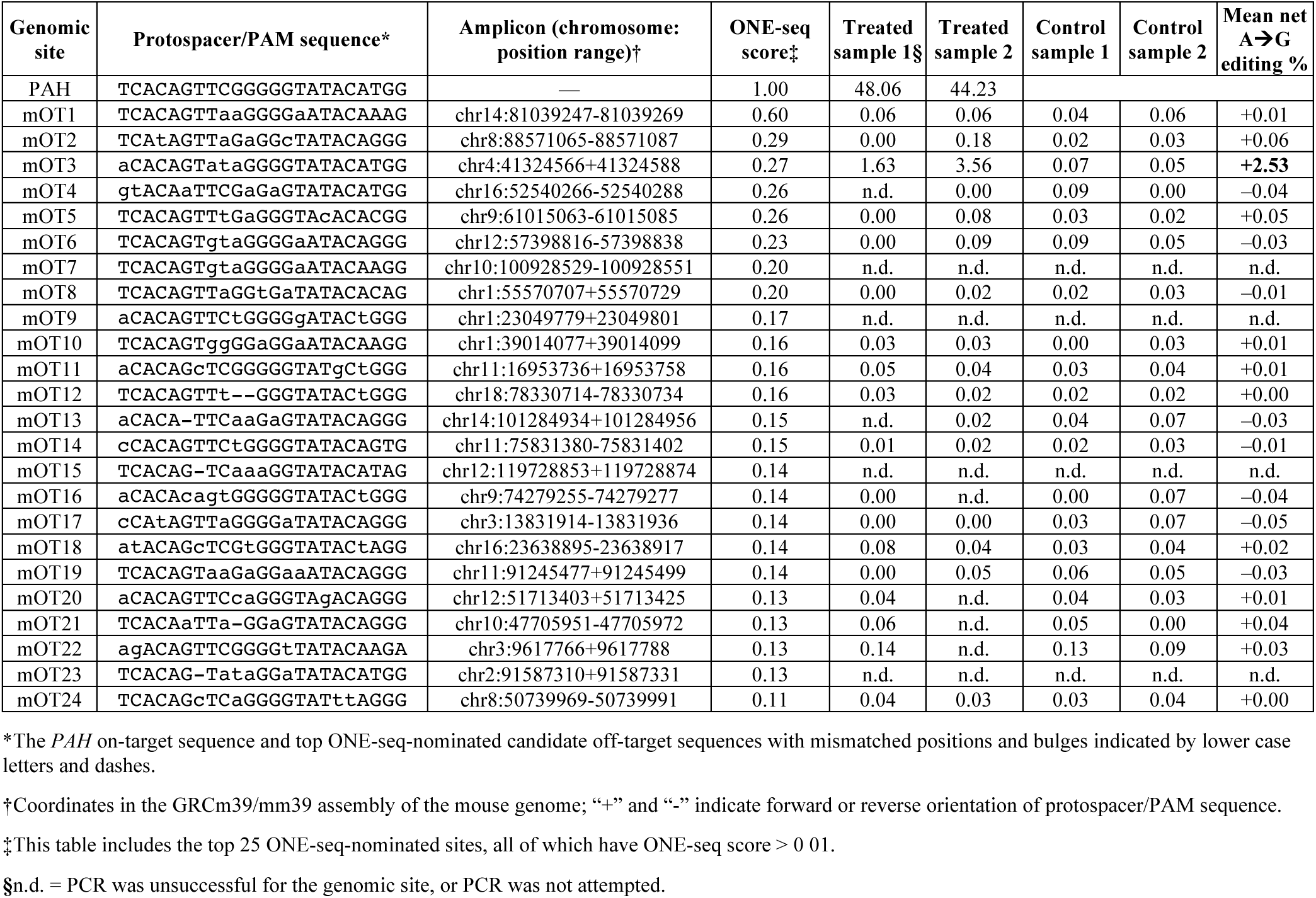
Assessment of off-target editing with ABE8.8/PAH1 with LNP delivery in the liver in humanized *PAH* P281L mice via targeted amplicon sequencing.

**Supplementary Table 3.**
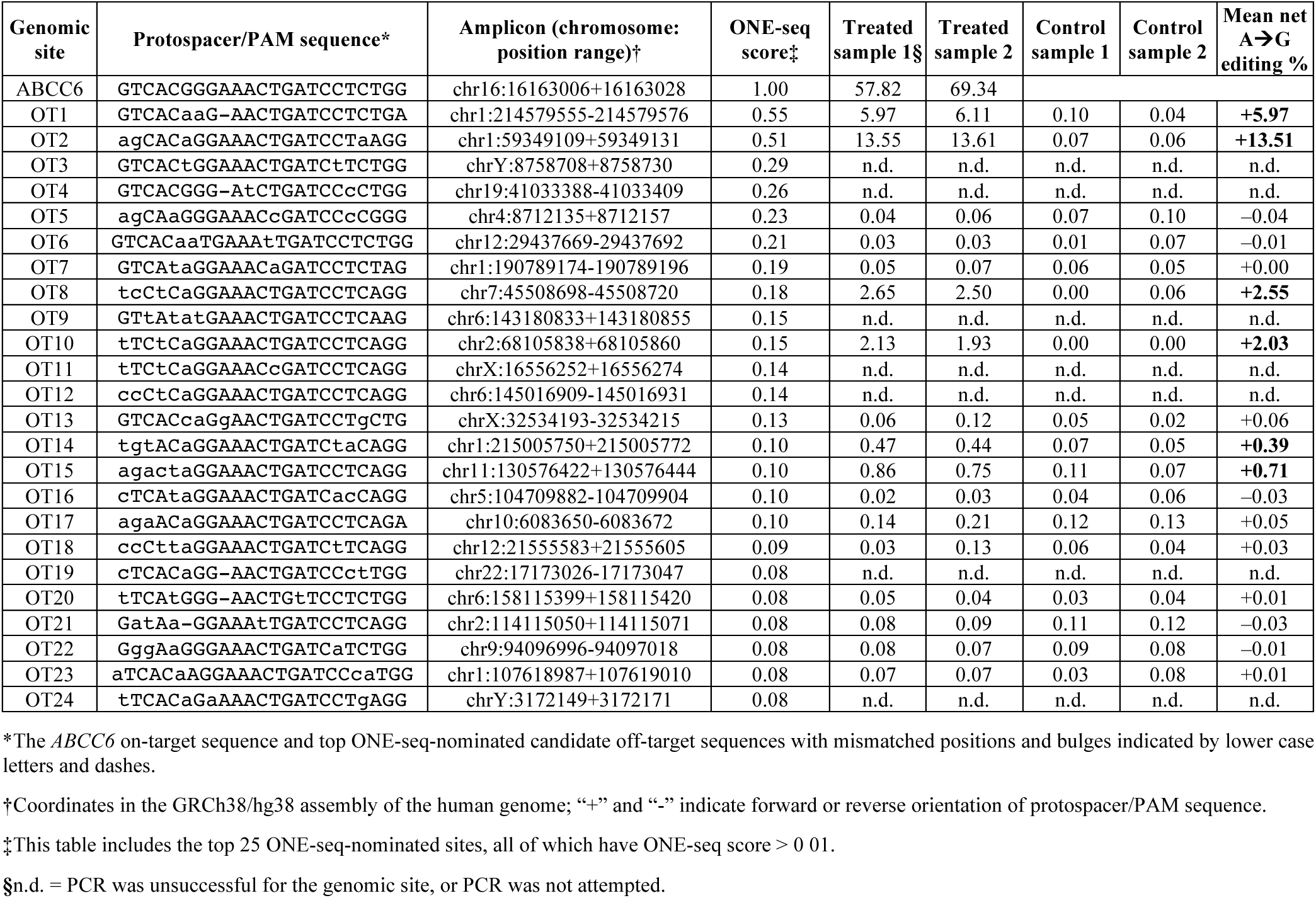
Assessment of off-target editing with ABE8.8/PXE1 with plasmid delivery in HuH-7 cells via targeted amplicon sequencing.

**Supplementary Table 4.**
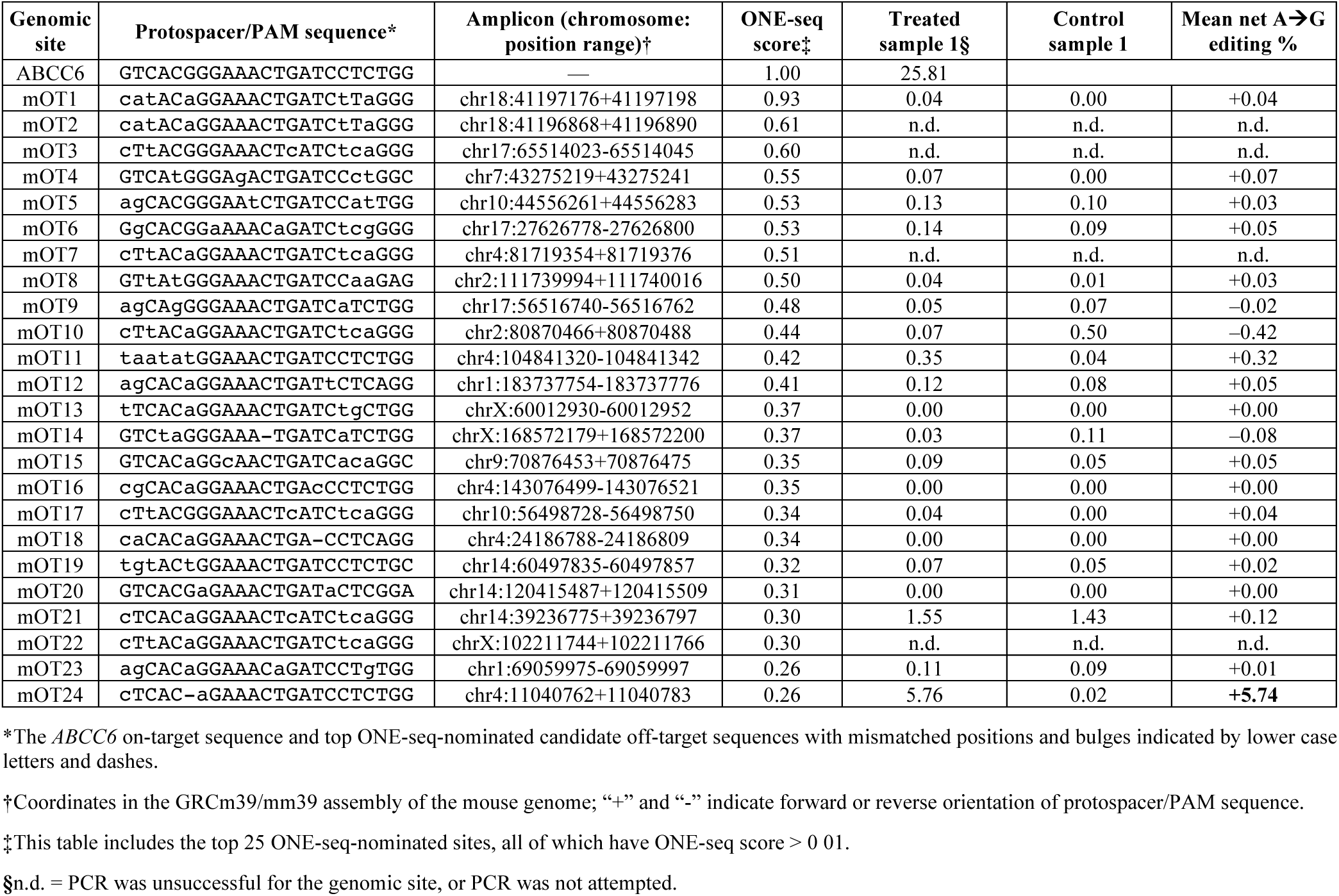
Assessment of off-target editing with ABE8.8/PXE1 with LNP delivery in the liver in humanized *ABCC6* R1164X mice via targeted amplicon sequencing.

**Supplementary Table 5.** Mean cytokine and chemokine levels in age-matched wild-type mice following LNP treatment or vehicle treatment.

| Time | IFN-γ | CXCL-1 | TNF-α | MCP-1 | IL-12p70 | RANTES | IL-1β | IP-10 | GM-CSF | IL-10 | IFN-β | IFN-α | IL-6 |
| --- | --- | --- | --- | --- | --- | --- | --- | --- | --- | --- | --- | --- | --- |
| **Treatment with phosphate-buffered saline (PBS vehicle), *N* = 4 animals per group** |  |  |  |  |  |  |  |  |  |  |  |  |  |
| **baseline** | 0.28 | 62.08 | 3.56 | 21.56 | 0.00 | 72.47 | 0.61 | 376.28 | 1.00 | 22.82 | 0.00 | 1.42 | 16.28 |
| **4 hours** | 0.00 | 90.95 | 3.62 | 51.41 | 0.00 | 32.38 | 0.72 | 288.18 | 0.00 | 0.00 | 0.00 | 0.00 | 398.51 |
| **24 hours** | 0.34 | 28.95 | 3.28 | 25.77 | 0.00 | 93.29 | 2.00 | 498.90 | 2.50 | 3.72 | 0.00 | 0.61 | 126.77 |
| **Treatment with ABE8.8/PXE1 LNPs, 2.5 mg/kg, *N* = 4 animals per group** |  |  |  |  |  |  |  |  |  |  |  |  |  |
| **baseline** | 0.09 | 38.55 | 3.37 | 0.00 | 0.00 | 30.05 | 1.31 | 346.69 | 0.00 | 5.06 | 0.00 | 0.00 | 3.92 |
| **4 hours** | 20.10 | 334.44 | 117.68 | 2189.41 | 0.72 | 243.91 | 57.32 | 13436.10 | 9.64 | 21.68 | 341.12 | 985.17 | 868.78 |
| **24 hours** | 0.21 | 27.44 | 5.84 | 94.07 | 0.00 | 126.71 | 5.50 | 1146.90 | 0.00 | 0.48 | 0.65 | 222.65 | 37.88 |
| **Treatment with ABE8.8/PXE1_hyb18 LNPs, 2.5 mg/kg, *N* = 4 animals per group** |  |  |  |  |  |  |  |  |  |  |  |  |  |
| **baseline** | 1.49 | 50.38 | 5.21 | 4.76 | 0.00 | 31.27 | 6.61 | 378.33 | 0.00 | 1.18 | 0.00 | 38.02 | 10.95 |
| **4 hours** | 4.63 | 405.06 | 57.88 | 689.26 | 0.35 | 91.57 | 25.67 | 5704.49 | 1.07 | 9.46 | 15.32 | 204.14 | 600.73 |
| **24 hours** | 5.70 | 38.33 | 13.49 | 54.62 | 0.21 | 77.39 | 15.20 | 1029.71 | 0.00 | 2.39 | 0.00 | 147.83 | 48.65 |

All cytokine/chemokine measurements are in units of pg/mL.

**Supplementary Table 6.**
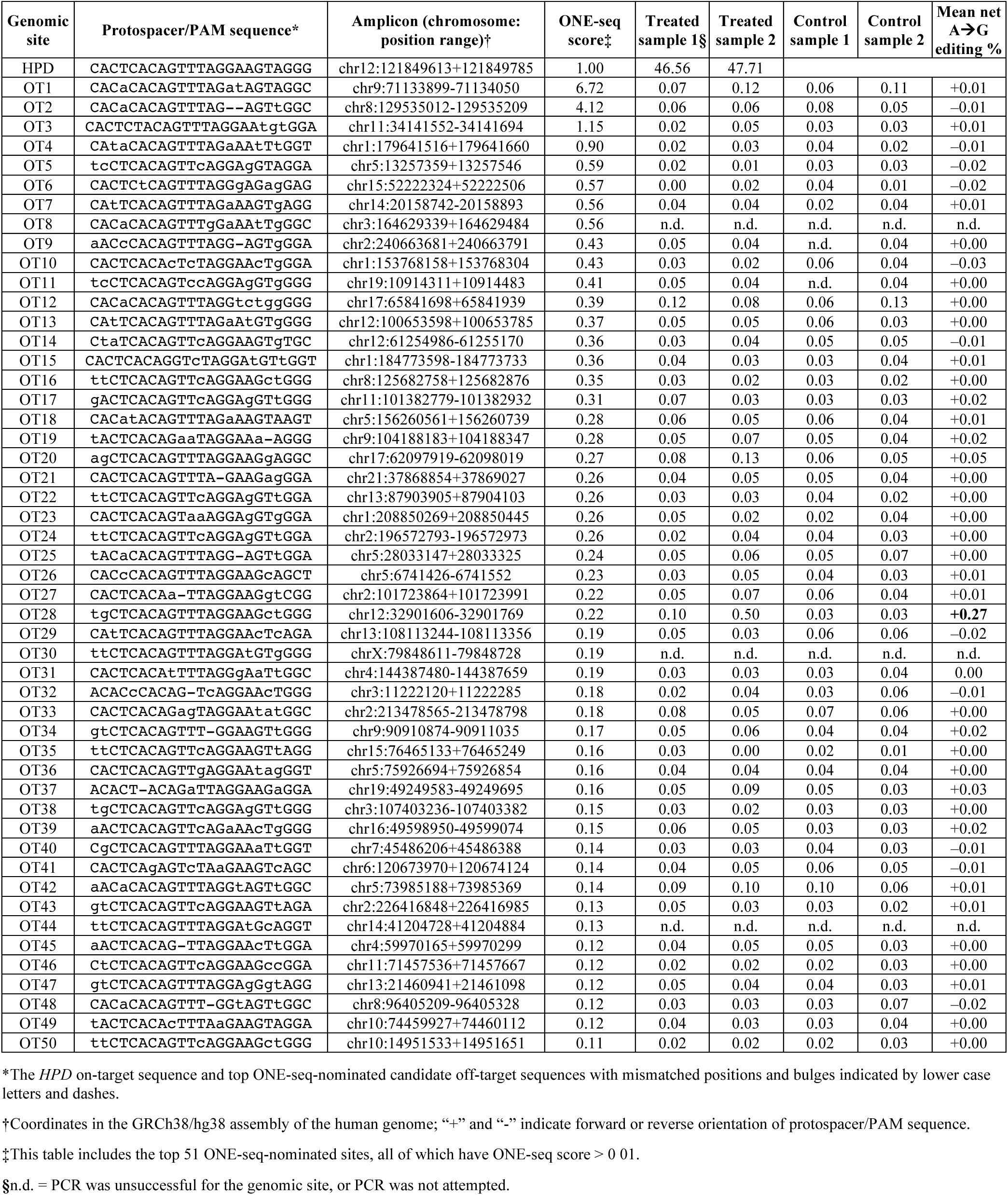
Assessment of off-target editing with ABE8.8/HPD3 with plasmid delivery in HuH-7 cells via targeted amplicon sequencing.

**Supplementary Table 7.**
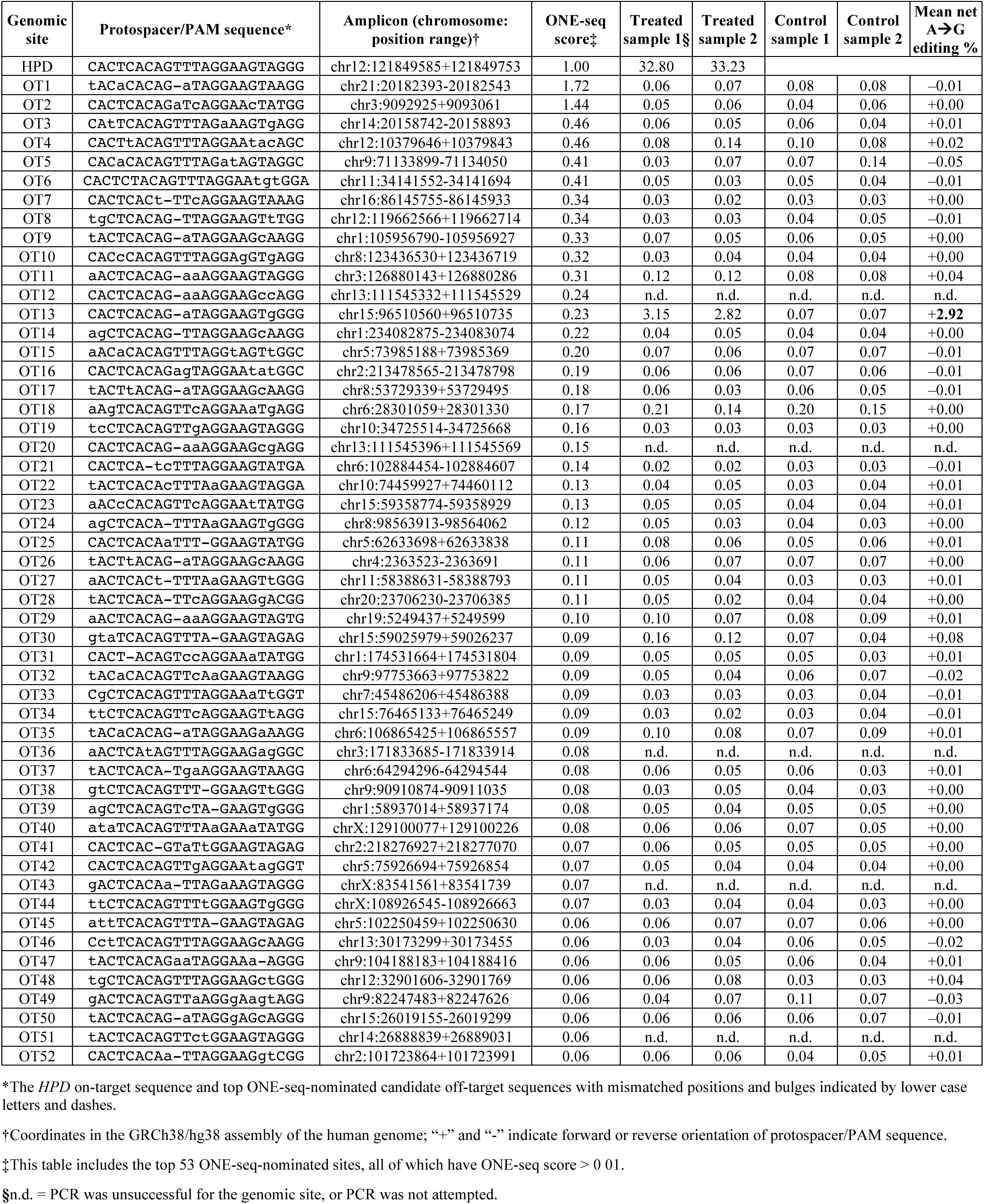
Assessment of off-target editing with ABE8.8/HPD4 with plasmid delivery in HuH-7 cells via targeted amplicon sequencing.

**Supplementary Table 8.**
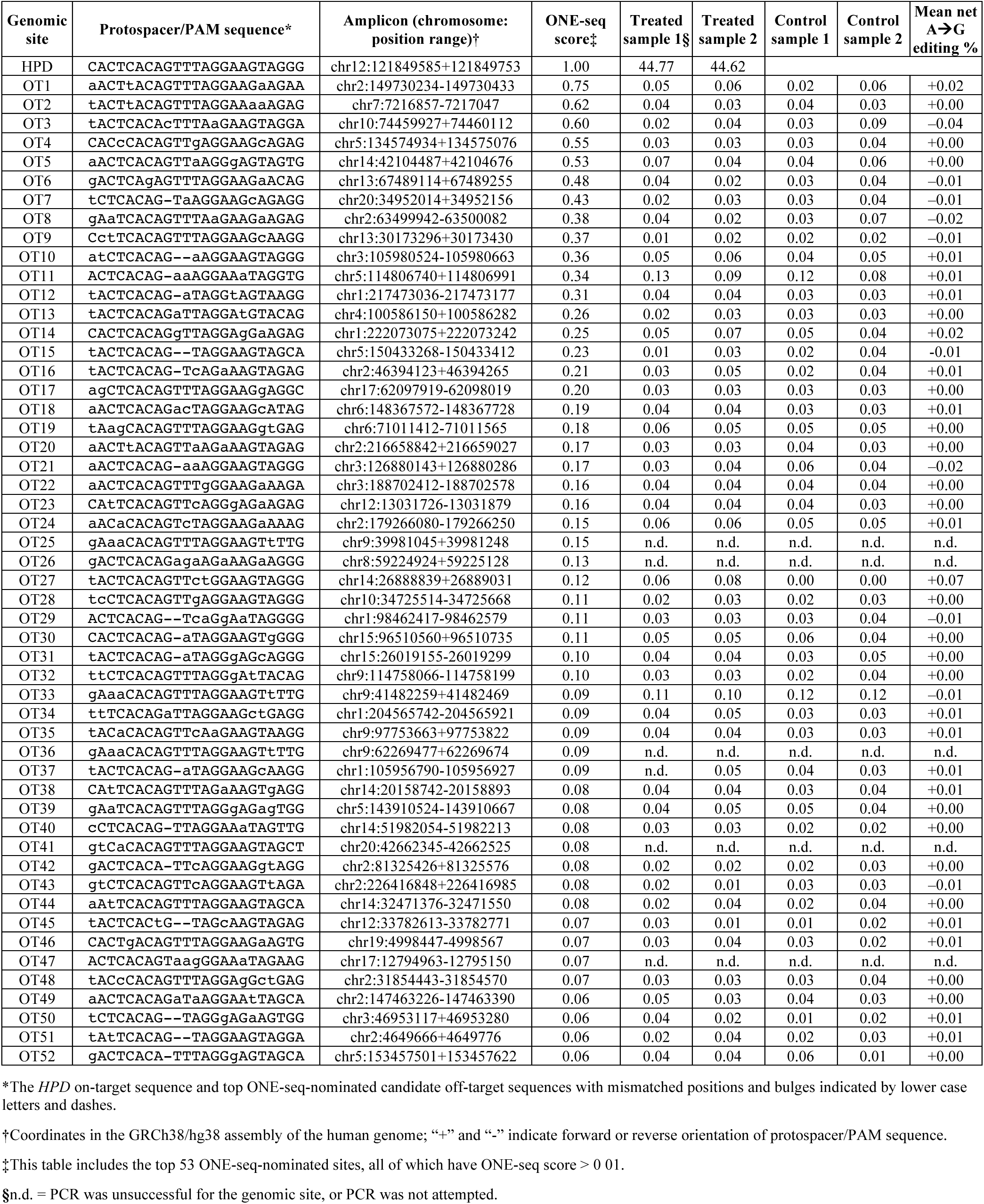
Assessment of off-target editing with ABE8.8/HPD5 with plasmid delivery in HuH-7 cells via targeted amplicon sequencing.

**Supplementary Table 9.**
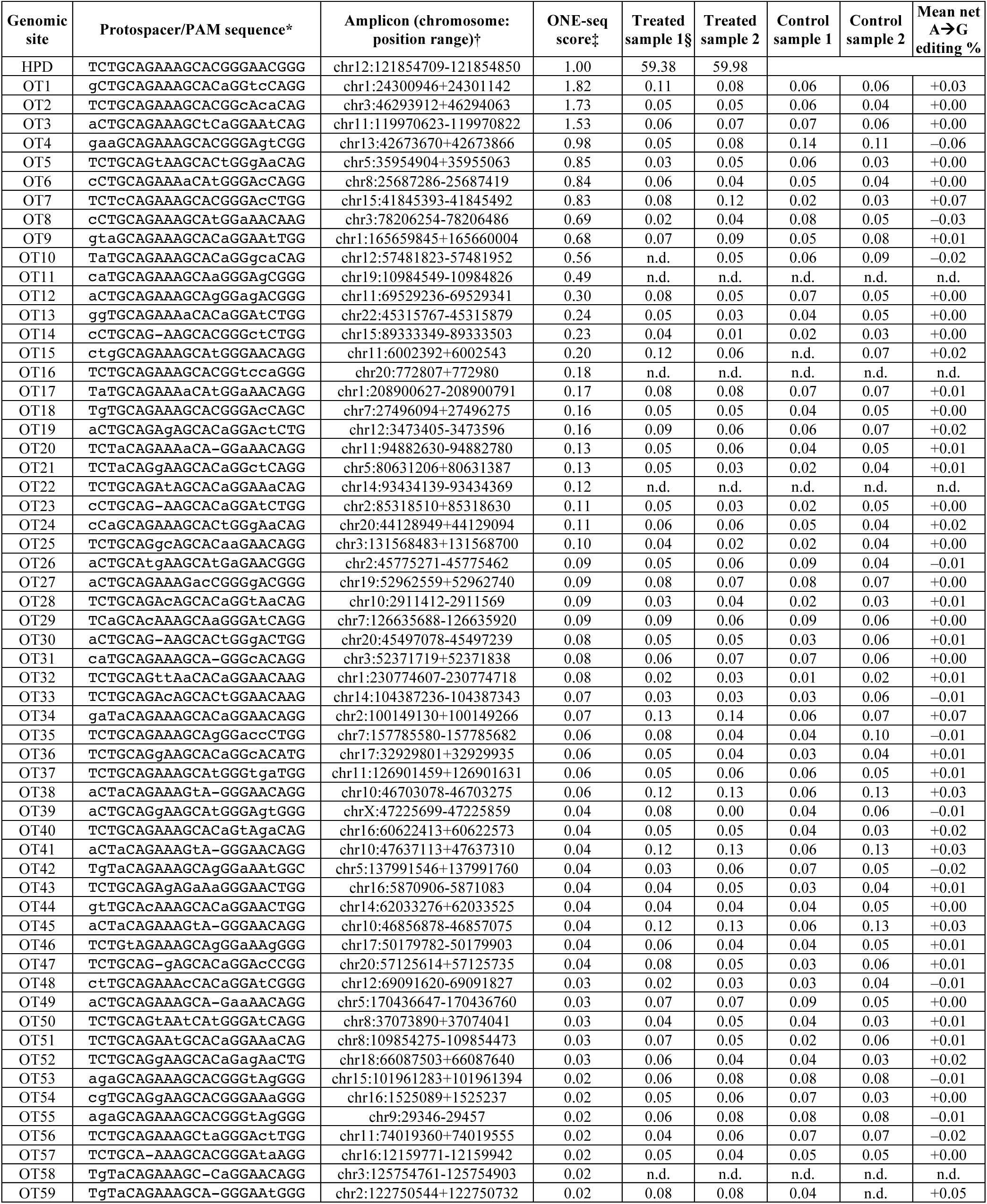
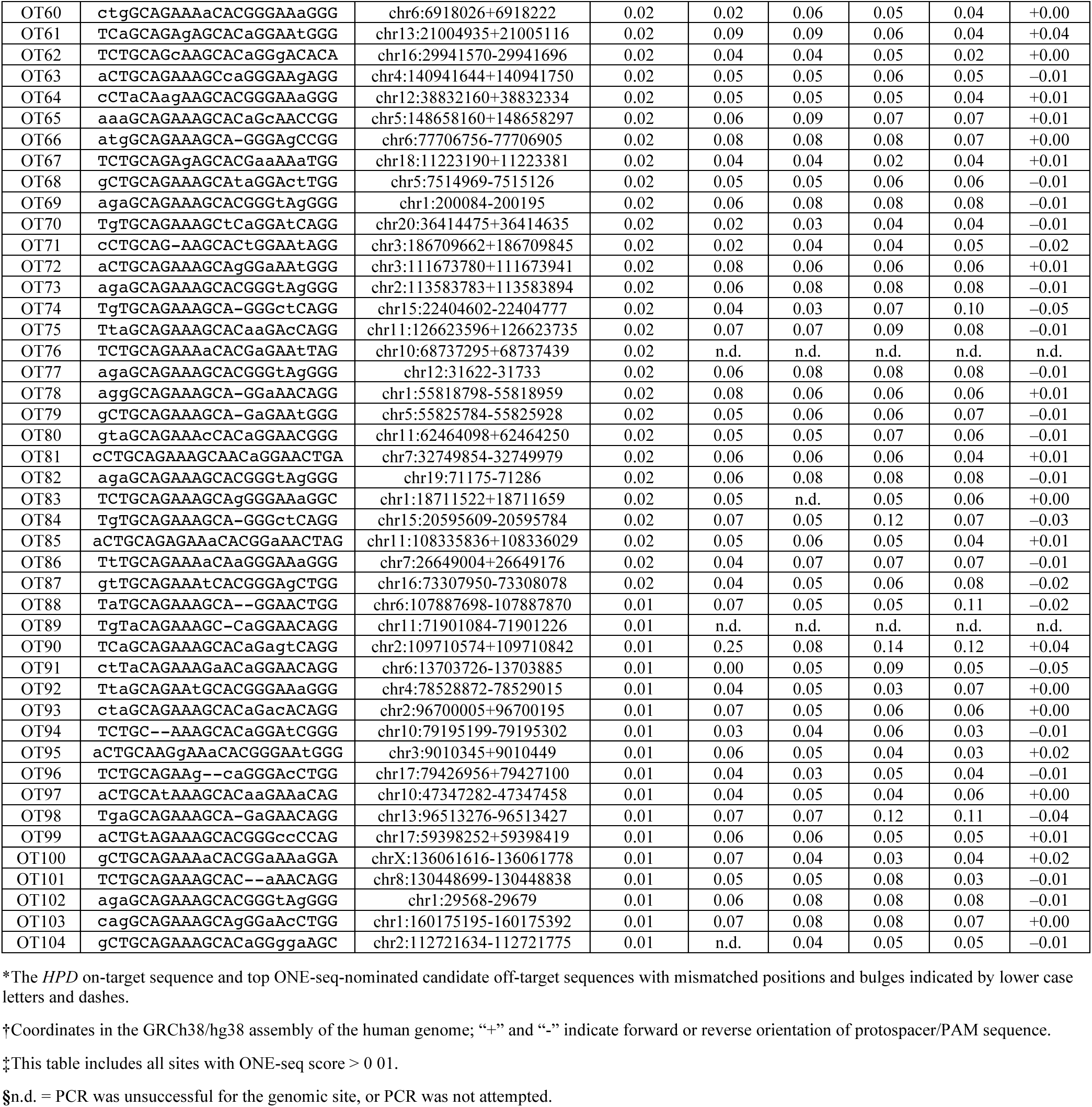
Assessment of off-target editing with ABE8.8/HPD20 with plasmid delivery in HuH-7 cells via targeted amplicon sequencing.

**Supplementary Table 10.**
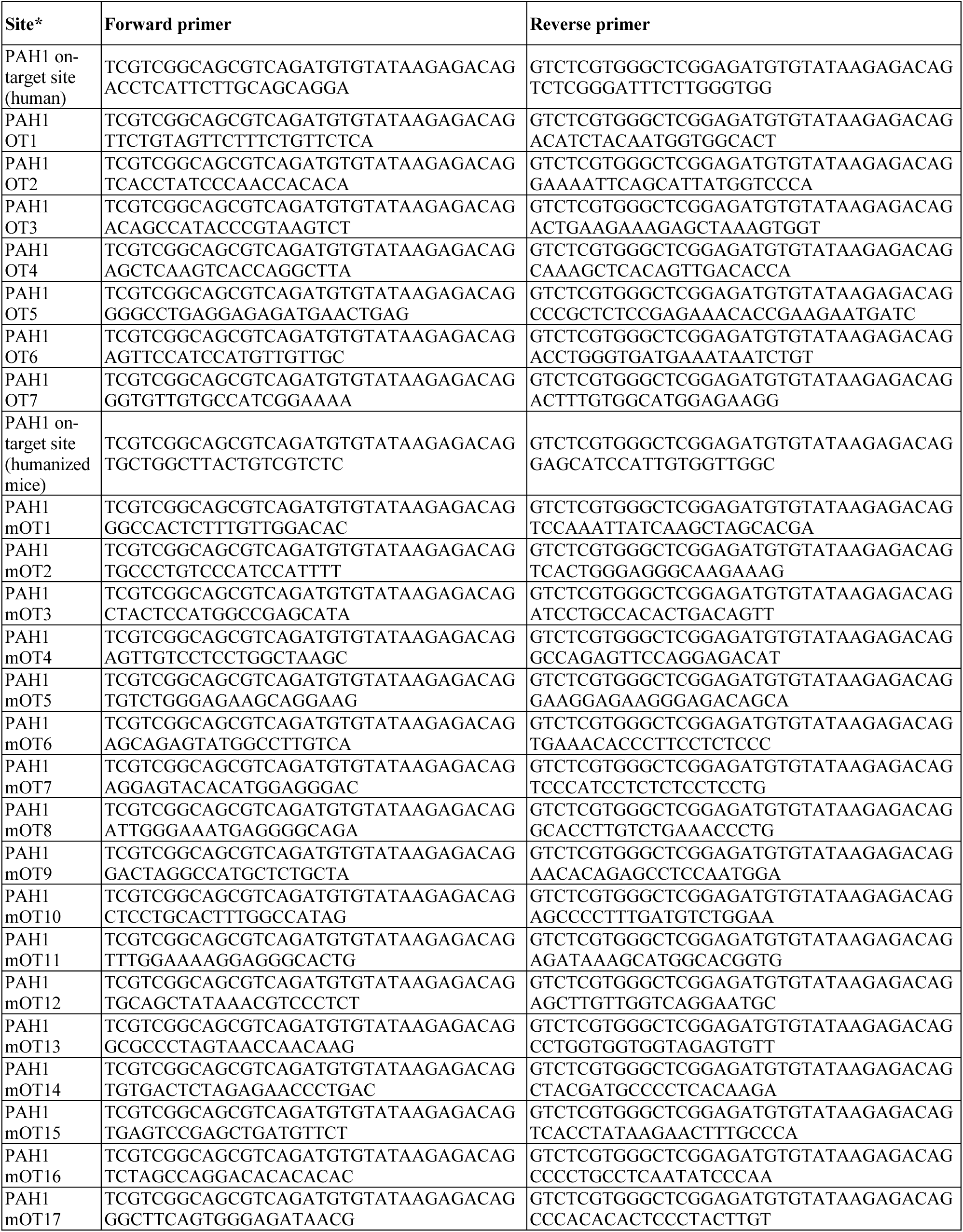
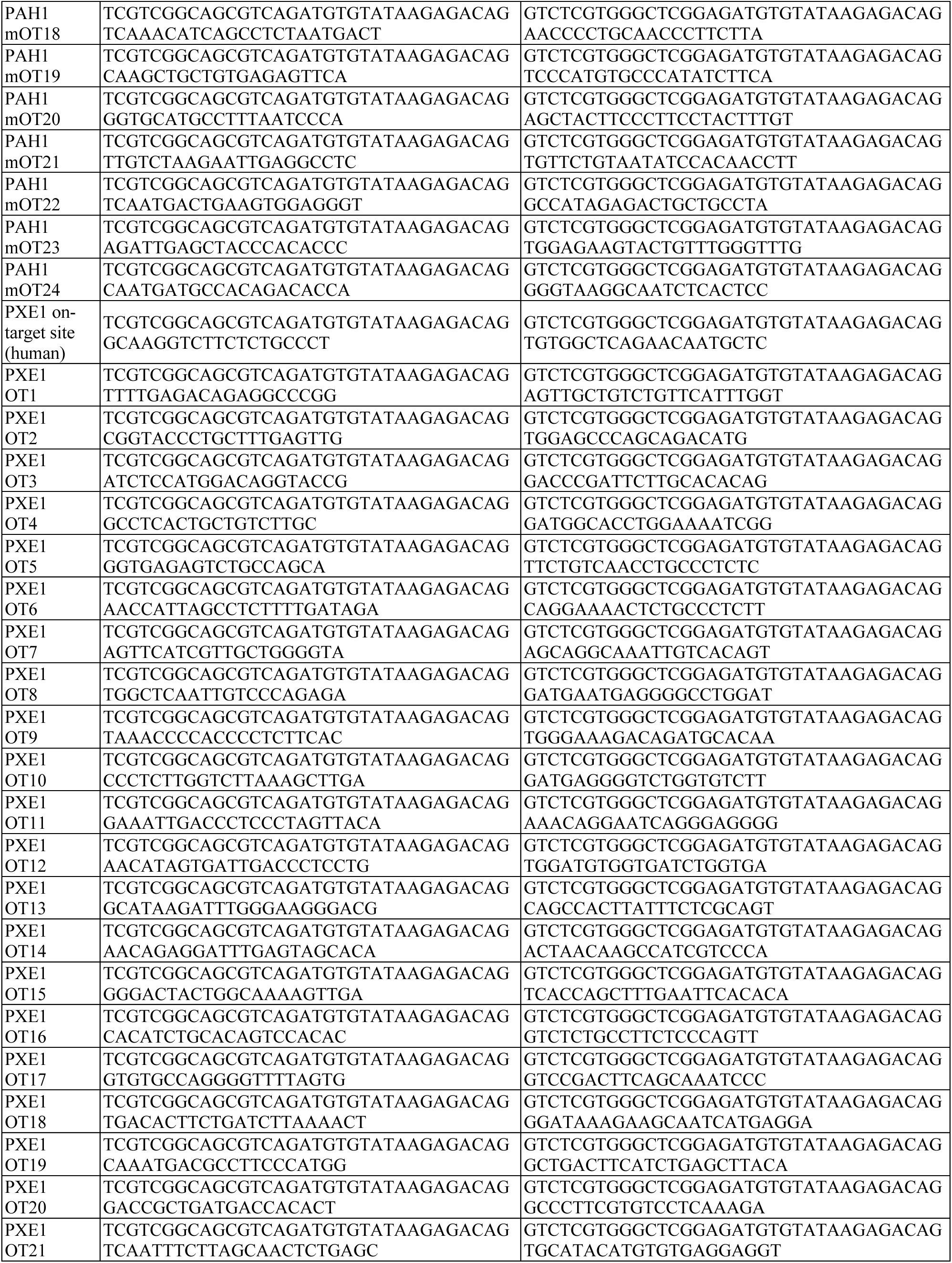
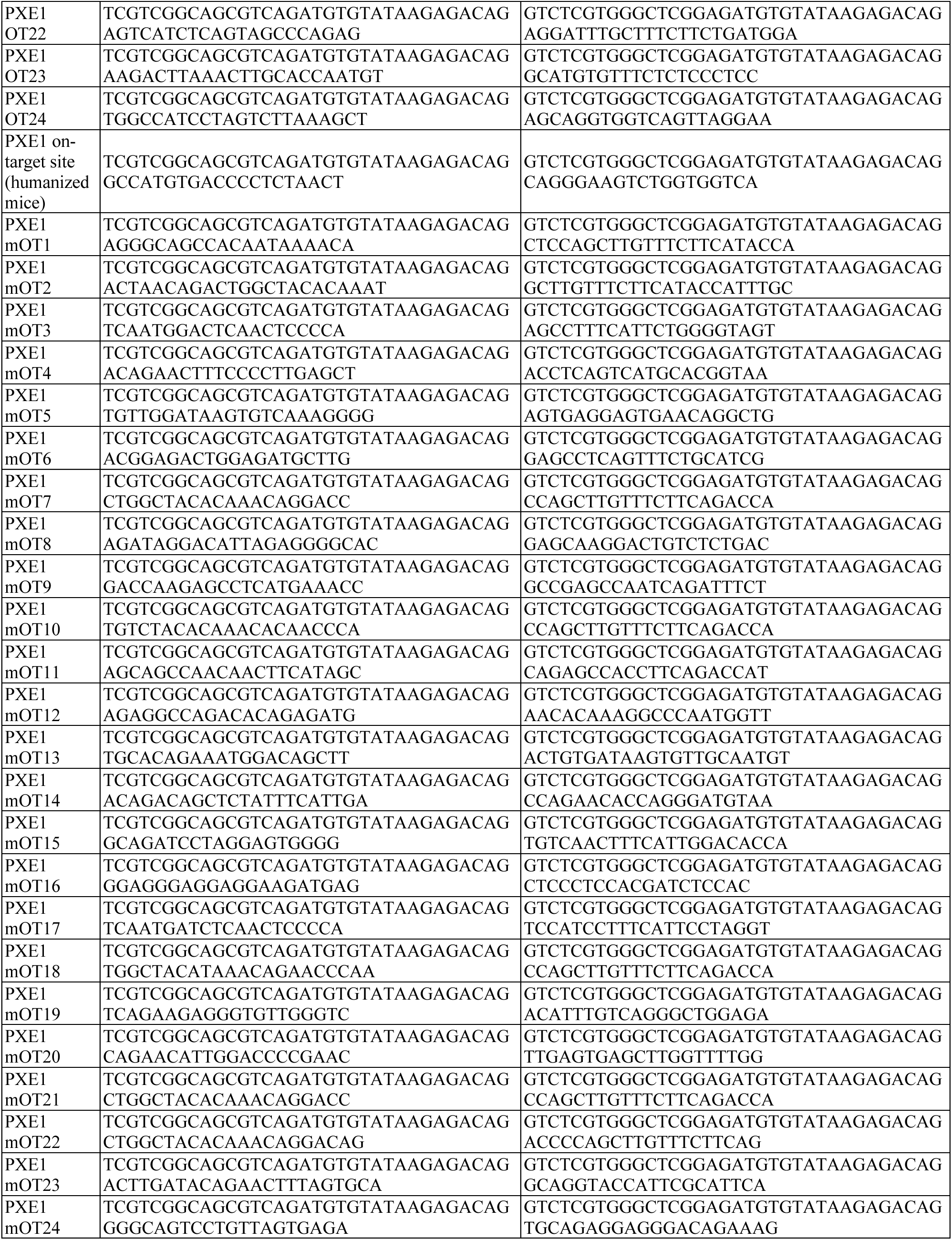
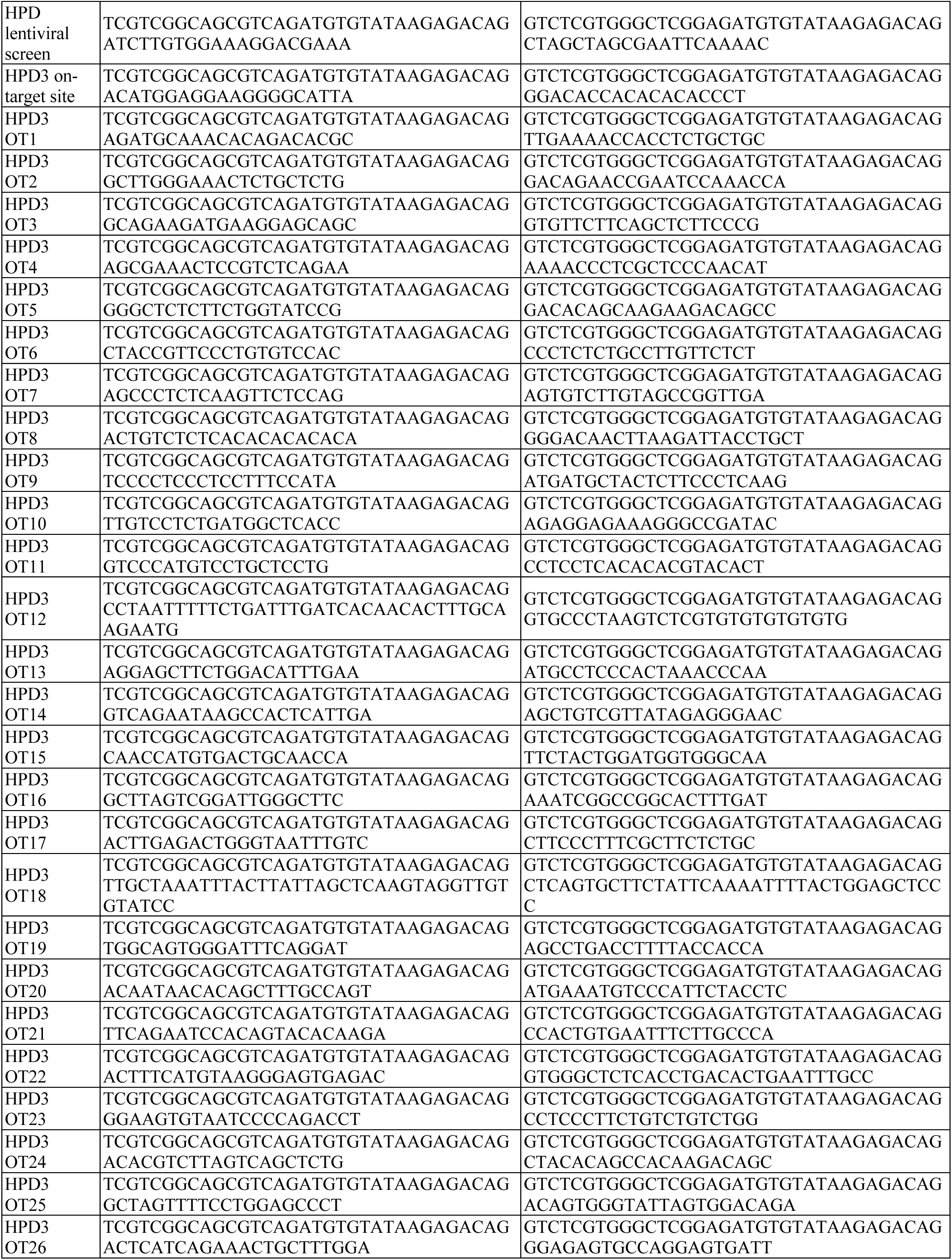
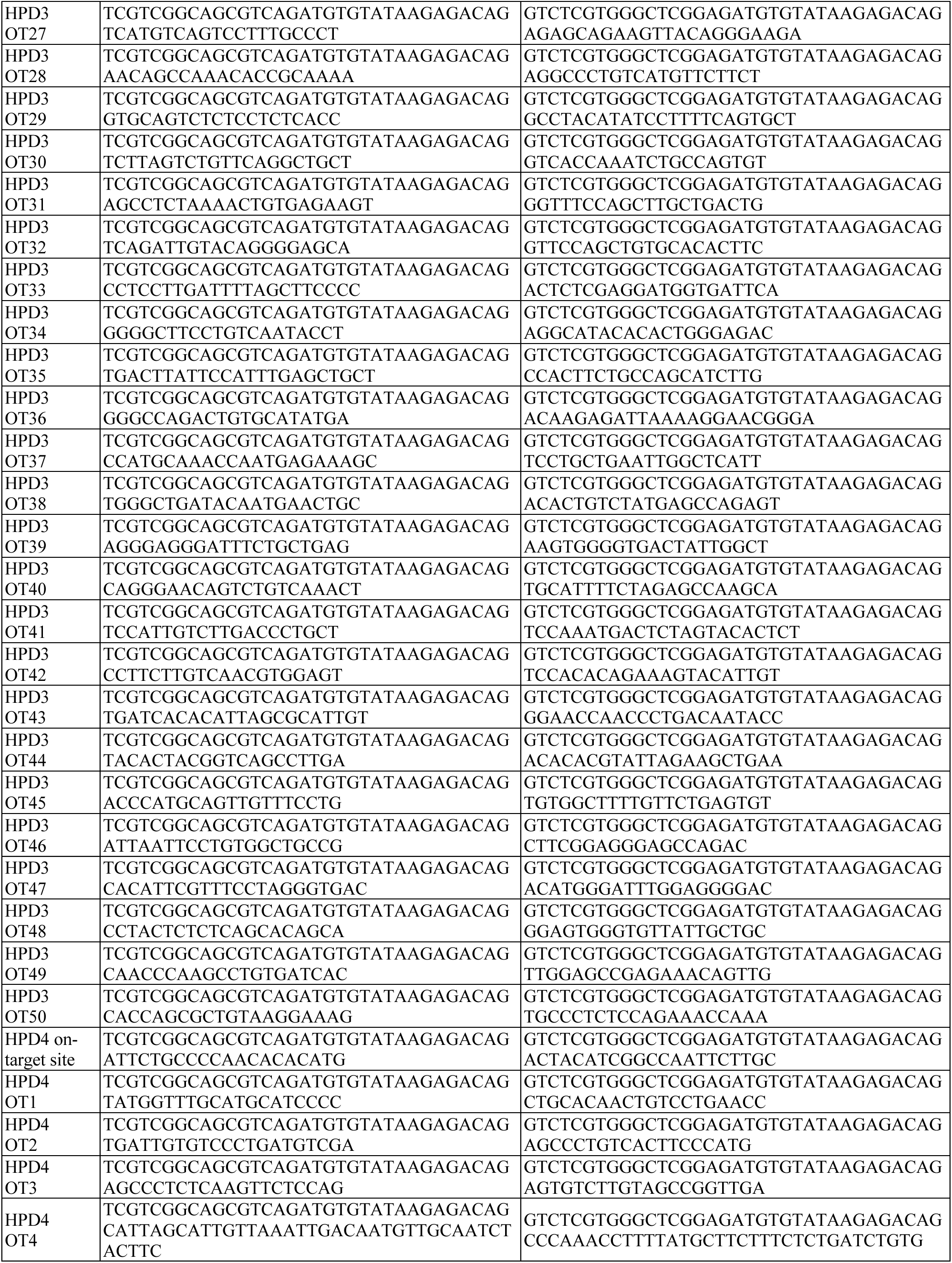
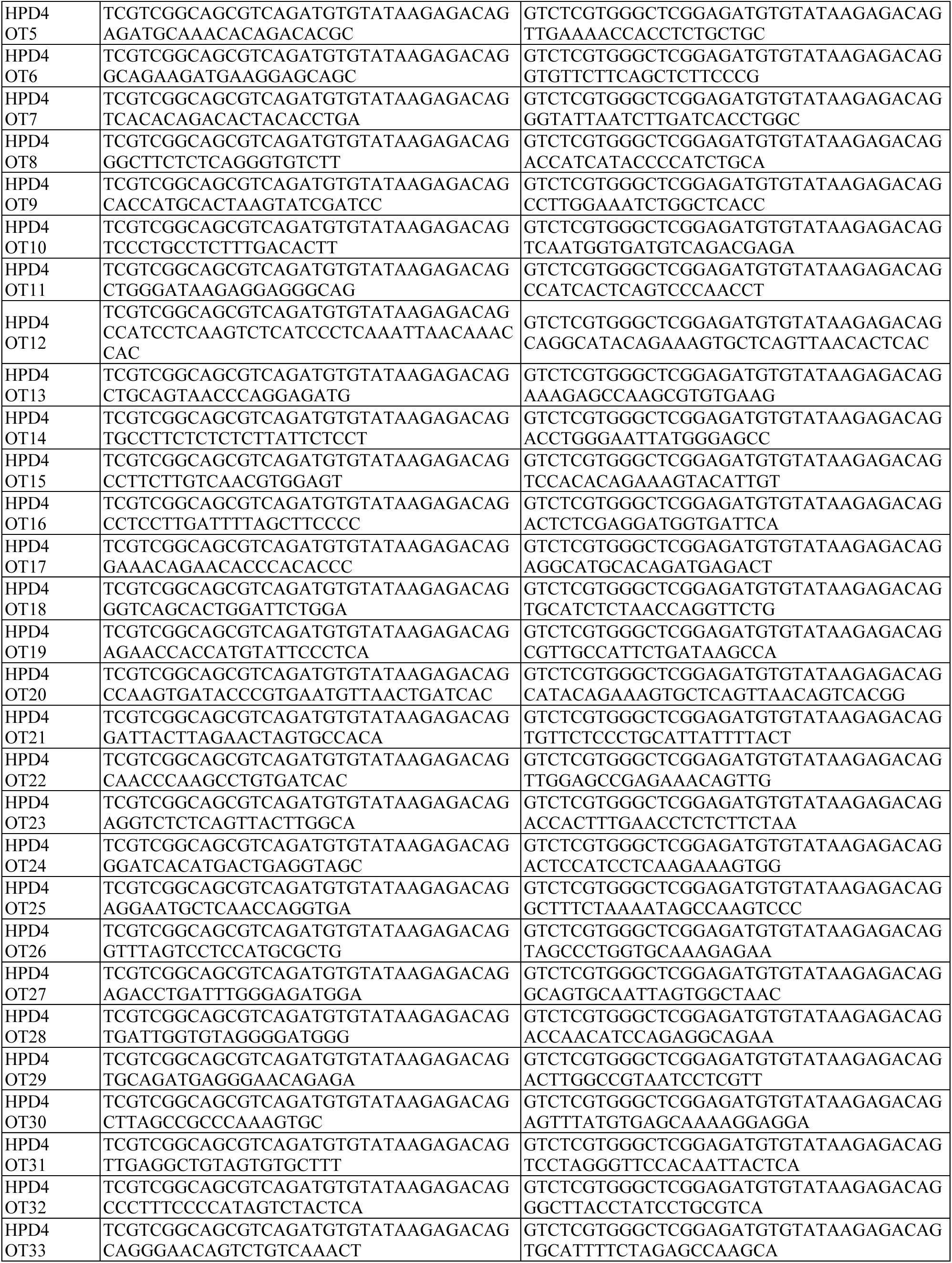
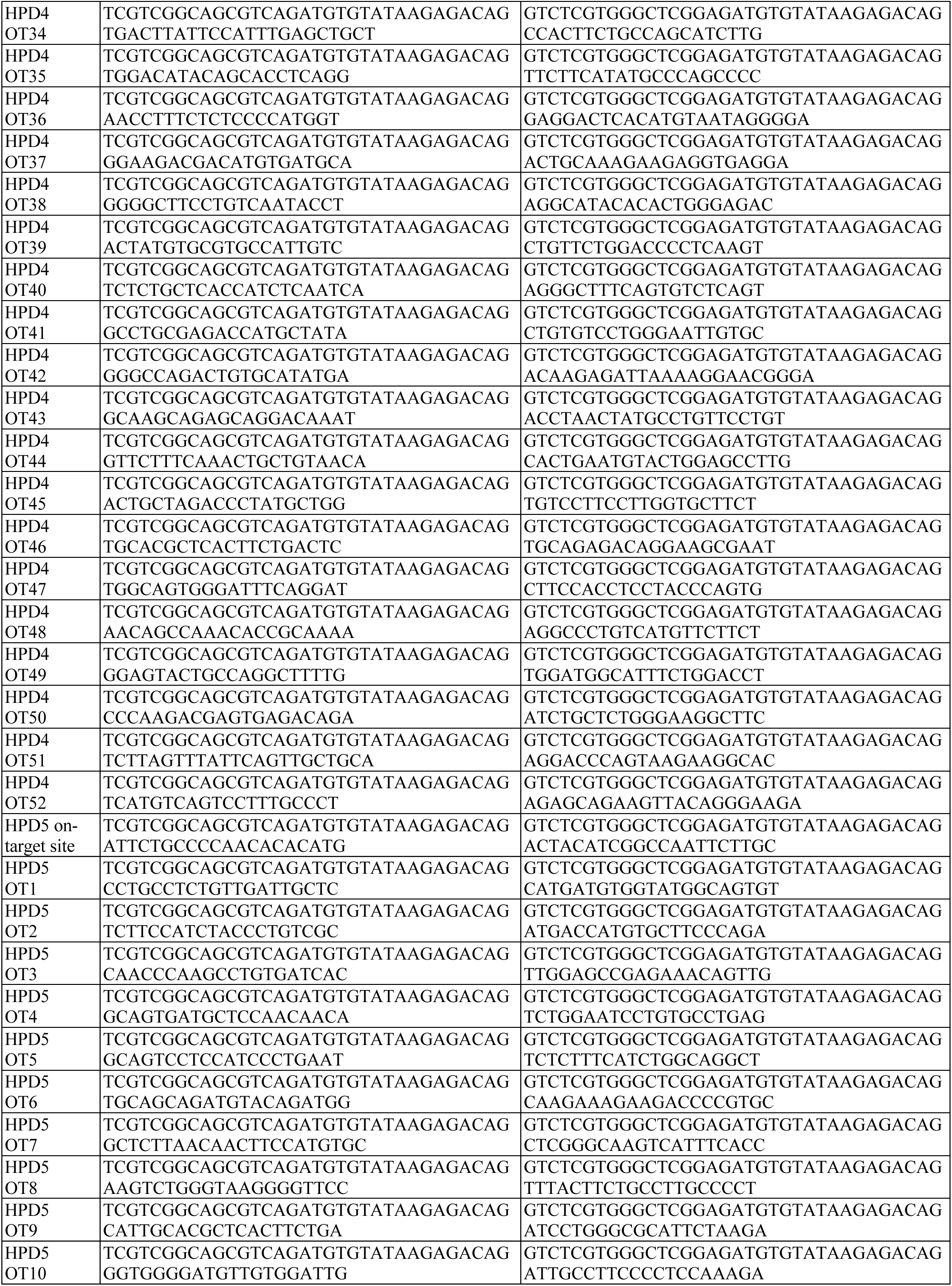
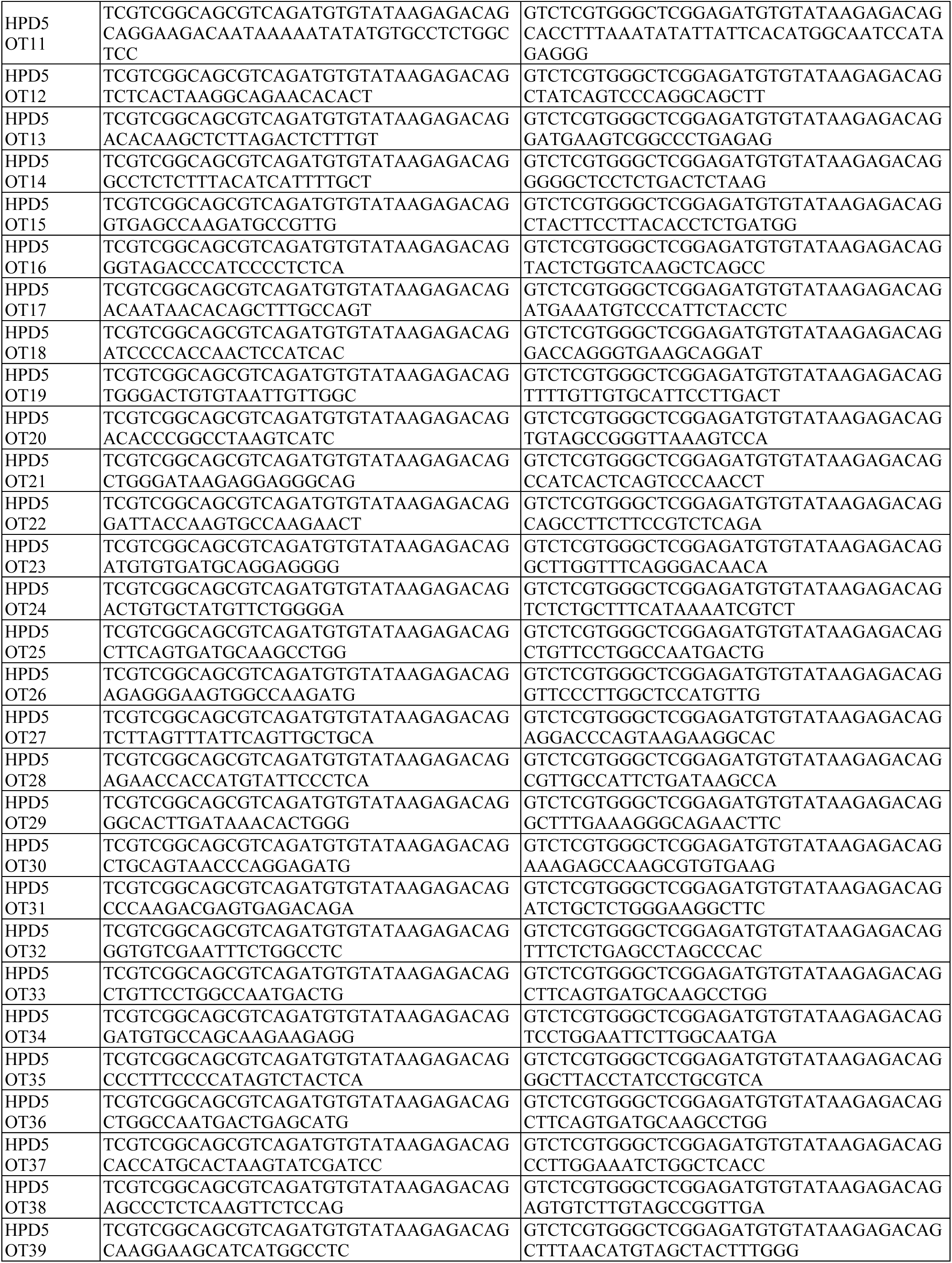
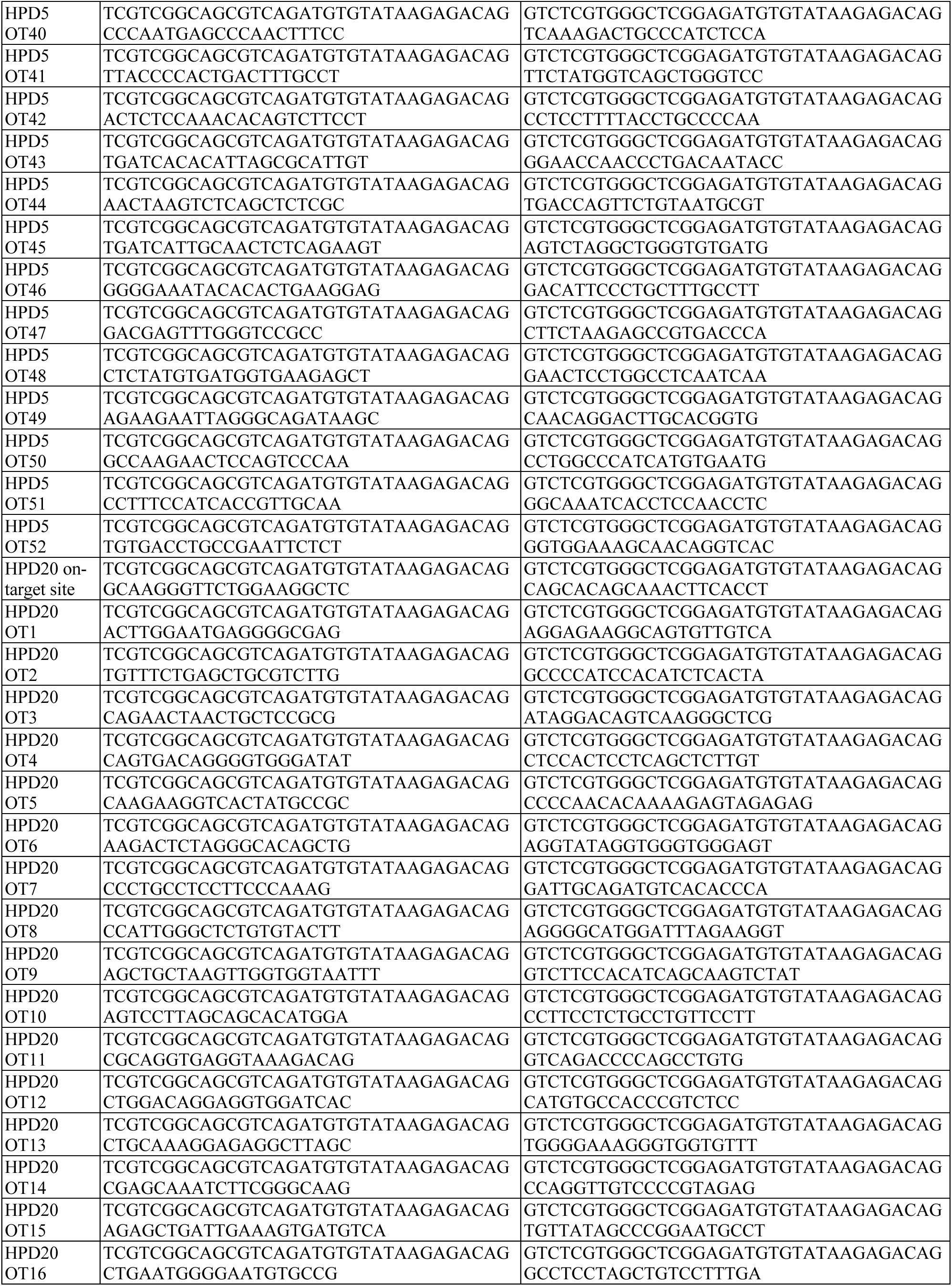
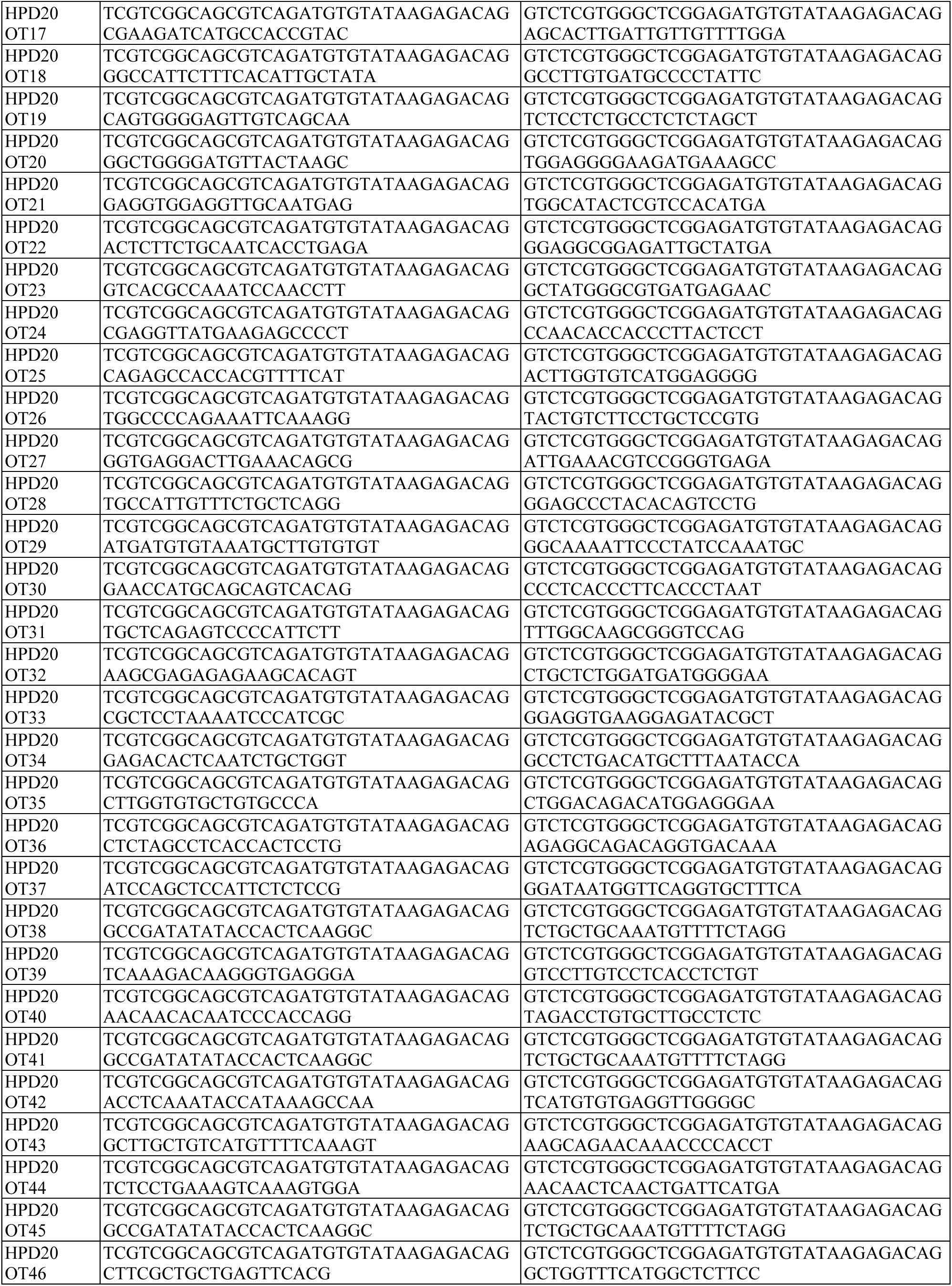
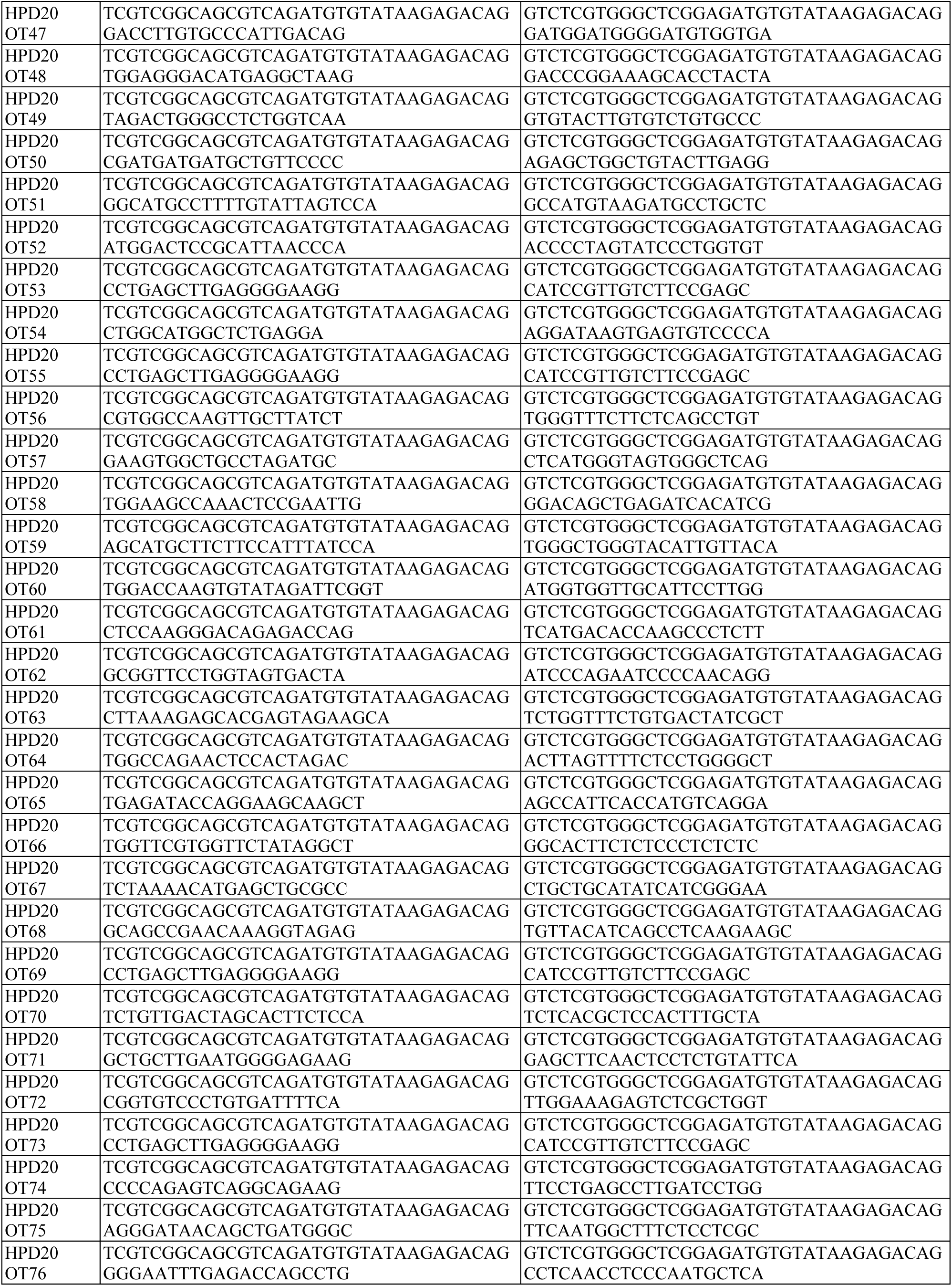
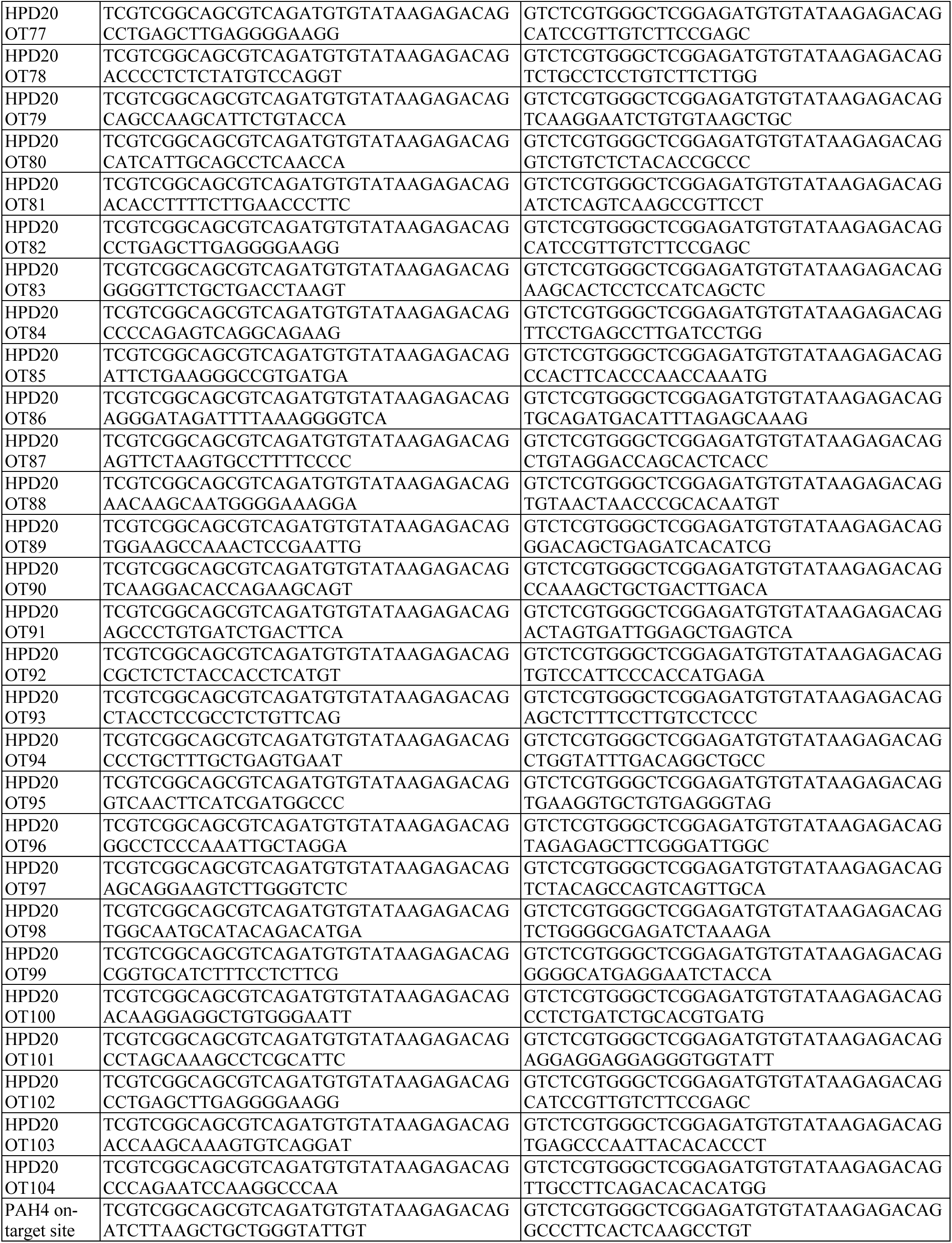
PCR primers for next-generation sequencing.

